# Memory trace superimposition impairs recall in a mouse model of AD

**DOI:** 10.1101/467043

**Authors:** Stefanie Poll, Lena C. Schmid, Julia Steffen, Jens Wagner, Boris Schmidt, Walker S. Jackson, Susanne Schoch, Dan Ehninger, Martin Fuhrmann

**Affiliations:** Neuroimmunology and Imaging Group, German Center for Neurodegenerative Diseases (DZNE), Bonn, 53127, Germany.; Clemens-Schöpf-Institute, Technical University of Darmstadt, 64289, Germany; Selective vulnerability of Neurodegenerative Diseases, German Center for Neurodegenerative Diseases (DZNE), Bonn, 53127, Germany; Institute of Neuropathology, University of Bonn, 53127, Germany; Molecular and Cellular Cognition Group, German Center for Neurodegenerative Diseases (DZNE), Bonn, 53127, Germany

**Keywords:** CA1, memory trace, engram, Alzheimer’s disease, IEGs, fosGFP, *c-fos*, novelty

## Abstract

Learning and memory processes depend on the hippocampus and are impaired in Alzheimer’s disease (AD). Active neuronal ensembles form an engram by encoding information during learning. Their reactivation is required for memory recall. However, it remains unresolved whether the engram in CA1 principal neurons is impaired under AD-like conditions. We used two-photon *in vivo* imaging to visualize the expression of the immediate early gene *c-fos* within CA1 neurons during contextual fear conditioning and retrieval. Surprisingly, we identified engrams in wild-type mice and in the mouse model of AD indicating intact memory formation. However, under AD-like conditions engrams were superimposed by a high number of newly recruited fosGFP^+^ neurons during memory recall. This superimposition resembled the network configuration of wild-type mice exposed to a novel context. Artificial superimposition of the memory trace during recall in wild-type mice was sufficient to induce memory impairment. Thus, we propose superimposition of the CA1 memory trace as a mechanism for memory impairment in a mouse model of AD.

**Highlights:**

- Decreased fosGFP expression in direct vicinity to amyloid-β plaques
- Intact engram in CA1 of APP/PS1 mice
- Impurity of the retrieval network in CA1 is sufficient to impair memory recall

Poll *et al*. present a novel mechanism for memory impairment in a mouse model of AD. The potential memory trace was found intact in the CA1 region of the hippocampus. However, excessive neuronal activity during retrieval, was superimposing the memory trace in a mouse model of AD.

## Introduction

Memories constitute our personality and evolve throughout life. Many theories exist regarding the formation, storage and recall of memories, but we are still far from completely understanding how the brain encodes and stores memories on the synaptic, cellular and systems level. Theories agree that memory traces are represented and evolve in hippocampal and prefrontal cortical networks (Nadel and Moscovitch, 1997; Richards and Frankland, 2013). An evolution of a memory trace on the cellular and systems level is proposed, meaning that overlapping, but different cells and systems like the hippocampus and prefrontal cortex are involved in memory consolidation in a time-dependent manner (Mankin, 2012). Moreover, several studies support the hypothesis that new learning does not occur in a “tabula rasa” (McKenzie and Eichenbaum, 2011), but rather has to be integrated within pre-existing memories (Frankland and Bontempi, 2005; Nadel and Moscovitch, 1997; Yiu et al., 2014). There have been significant advances in the knowledge about reactivation of memories and the subsequent distribution from hippocampal to prefrontal networks (Czajkowski et al., 2014; Gdalyahu et al., 2013; Guzowski et al., 1999; Han et al., 2009; Reijmers et al., 2007; Richards et al., 2014; Tayler et al., 2013; Trachtenberg et al., 2002). Especially the development of optogenetics enabled the identification of engram cells that are necessary to retrieve a memory and sufficient to induce artificial retrieval (Cowansage et al., 2014; Ramirez et al., 2013). AD is characterized by memory impairment (Palop and Mucke, 2010b), a feature that was successfully recapitulated in animal models of the disease (Jankowsky et al., 2004). The neuronal network dysfunction hypothesis argues that changes on the molecular, synaptic, cellular and circuit level ultimately lead to neuronal network dysfunction, that presumably causes the spontaneously occurring fluctuations in neurological functions of AD patients (Palop and Mucke, 2010a). Indeed, network dysfunction as indicated by epileptiform activity has been found to be associated with AD and mouse models of AD (Palop et al., 2007; Palop and Mucke, 2010b). On the cellular level hyperactive neurons have been identified in the proximity of amyloid- (A plaques in the cortex and hippocampus of an AD mouse model presumably representing one reason for network dysfunction (Busche et al., 2012; Busche et al., 2008). In the visual cortex expression of the immediate early gene (IEG) *Arc*, an indicator of neuronal activity, was increased in proximity to A plaques, indicating hyperactivity of neurons under AD-like conditions. Furthermore, visual stimulation with a light-dark paradigm lead to disturbed *Arc* expression (Rudinskiy et al., 2012). In humans, hippocampal hyperactivity was reported to precede a profound loss of hippocampal activity, which is accompanied by clinical decline measured by the clinical dementia rating scale (CDR) (O’Brien et al., 2010). IEG expression as a correlate of neuronal activity has been used for the identification of memory traces in wild-type animals (Guzowski et al., 1999; Han et al., 2009; Ramirez et al., 2013; Reijmers et al., 2007), and recently the artificial activation of dentate gyrus neurons belonging to a memory trace rescued fear memory at early disease stages in a mouse model of AD (Roy et al., 2016). However, whether the composition of memory traces in the CA1, the major output region of the hippocampus, is affected under AD-like conditions and how the potential disturbance relates to memory impairment remains unknown.

We hypothesized that either encoding or recall of a memory was impaired under AD-like conditions, which would be reflected in the cellular composition of the memory traces in the hippocampal CA1 region. To test this hypothesis, we improved the classical two time-point memory trace analysis that compares IEG expression between memory encoding and retrieval. We measured IEG expression in the same dorsal CA1 neurons up to ten times during encoding and retrieval of a hippocampus-dependent contextual fear memory using repetitive two-photon *in vivo* imaging. We identified memory traces in healthy and diseased mice indicating intact memory encoding in the mouse model of AD. Furthermore, we demonstrate a novel mechanism for impaired memory retrieval under AD-like conditions: Superimposition of the memory trace by aberrantly active hippocampal CA1 neurons during recall.

## Results

### Decreased fosGFP expression in proximity to Aβ plaques *in vivo*

The IEG *c-fos* is a well-described marker for neuronal activity (Sagar et al., 1988; Schoenenberger et al., 2009). We utilized fosGFP mice crossbred to the APP/PS1 mouse model of AD, from here on referred to as wild-type and APP/PS1 mice (Barth et al., 2004; Jankowsky et al., 2004) (Supplementary Fig. 1A, B). These mice reliably expressed endogenous Fos in fosGFP-positive (fosGFP^+^) neurons (>96%; Supplementary Fig. 1C,D). Furthermore, fosGFP^+^ neurons in stratum pyramidale of the dorsal hippocampal CA1 region were putatively excitatory neurons, since we rarely (<1%) identified fosGFP^+^ inhibitory neurons (Supplementary Fig. 1E,F). To analyze the level of GFP expression in fosGFP^+^ neurons in dorsal hippocampus we installed a chronic cranial window and carried out two-photon *in vivo* imaging as previously described (Gu et al., 2014; Schmid et al., 2016) (Fig. 1AC). Since aberrant expression levels of IEGs have been identified close to A plaques (Busche et al., 2012; Busche et al., 2008; Rudinskiy et al., 2012), we asked whether the density of fosGFP^+^ neurons or fosGFP expression was likewise altered in our mouse model. The density of fosGFP^+^ neurons in APP/PS1 mice was similar to wild-type mice, and not altered in proximity to A plaques or randomly placed virtual A plaques in APP/PS1 and wild-type mice, respectively (Fig. 1D-F). However, fosGFP fluorescence intensity in hippocampal CA1 neurons of APP/PS1 mice was significantly decreased (Fig. 1G,H), an effect observed only in neurons residing closer than 50 µm to the next Aβ plaque (Fig. 1I). These data suggest decreased neuronal activity of CA1 neurons in proximity to Aβ plaques.

**Figure 1.**
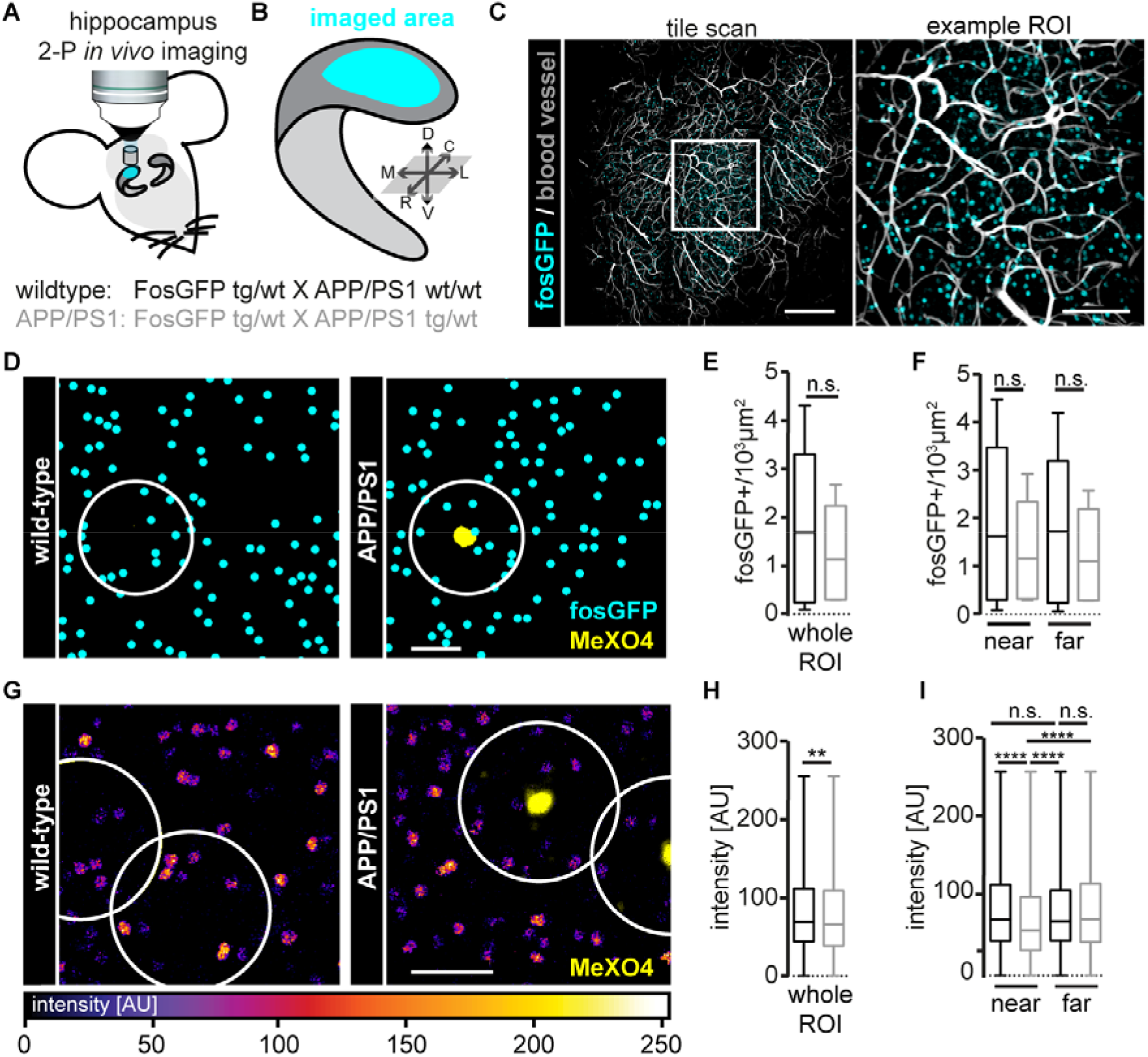
*In vivo* fosGFP expression of CA1 neurons is decreased in the vicinity of Aß plaques. **(A, B)** Schematic of a chronic cranial window preparation for hippocampal two-photon *in vivo* imaging (A) and imaged dorsal hippocampal region in blue (B). **(C)** Tile scan (left) and zoom (right, boxed region in left panel) displaying fosGFP^+^ neurons (cyan) in hippocampal CA1 area. Blood vessels (light grey) were visualized with a texas-red dextran tail vein injection. ROI, region of interest. **(D)** Exemplary images showing the density of fosGFP^+^ neurons around Methoxy-XO4 (MeXO4) stained Aß plaques and around randomly placed virtual Aß plaques of APP/PS1 and wild-type mice, respectively. **(E, F)** Density of fosGFP^+^ neurons irrespective of Aß-plaque distance (E), and in the proximity (<50 µm, near) or distant (>50 µm, far) to MeXO4 stained Aß plaques (F) comparing APP/PS1 and wild-type mice, respectively. **(G)** Exemplary images of fosGFP^+^ neurons’ fluorescence intensities in the hippocampus of wild-type and APP/PS1 mice. **(H, I)** Fluorescence intensity of fosGFP^+^ neurons in wild-type and APP/PS1 mice regardless of Aß-plaque distance (H), and in proximity (<50 µm, near) or distant (>50 µm, far) to MeXO4 stained Aß plaques (I). Data from n=8 wild-type (3385 fosGFP^+^ neurons) and n=6 APP/PS1 mice (2286 fosGFP^+^ neurons); (E) p=0.4342, unpaired t-test; (F) p for every comparison > 0.05, ordinary one-way ANOVA with Holm-Sidak’s correction for multiple comparisons. (H) **p=0.0032, Mann-Whitney test; (I) ****p < 0.0001, Kruskal-Wallis test with Dunn’s correction for multiple comparisons. (D, G) White circle, radius = 50 µm. Scale bars: (C) 250, 100 µm; (D, E) 50 µm. See also Figure S1.

### Longitudinal *in vivo* imaging revealed different subsets of fosGFP neurons

Since we aimed at identifying and comparing memory traces of wild-type and APP/PS1 mice by recording fosGFP expression throughout a hippocampus-dependent memory task, we first acquired fosGFP expression baseline changes, daily for a period of five days (Fig. 2). During this period, mice were only exposed to their home cage environment (HC) (Fig. 2A). After data acquisition, we generated binary images to visualize neurons that expressed fosGFP above or below threshold to define active and silent neurons, respectively (Supplementary Fig. 2A-C). Subsequently, neurons were categorized and color-coded according to their daily change in fosGFP expression: turning on (ON, green), switching off (OFF, magenta) or continuing expression (CON, blue) (Fig. 2B). This categorization revealed a large fraction (60%) of fosGFP^+^ neurons that continuously expressed fosGFP suggesting that these neurons constitute the “home cage” network (Fig. 2C). In addition, we identified equal fractions of ON and OFF neurons representing about 20%, each (Fig. 2C). The fractions of ON, OFF and CON neurons remained constant during the five-day baseline period, underscoring the stability of the neuronal network. APP/PS1 mice exhibited similar fractions of ON, OFF and CON neurons as wild-type mice, indicating an intact “home cage” network during the five-day baseline period (Fig. 2C-E). In addition to daily changes in fosGFP expression, we analyzed the expression duration of individual fosGFP^+^ neurons. Consistent with our previous results, we identified two major fractions in both, wild-type and APP/PS1 mice: fosGFP^+^ neurons with 1-day (~30%) and 5-day (~40%) expression duration (Supplementary Fig. 2D, E). We conclude that there exists an intermingled network state in hippocampal CA1, consisting of two neuronal populations: one continuously/repeatedly active subset of neurons, presumably representing stable contextual information and a variably active subset of neurons, able to adapt to changes in the environment. Furthermore, wild-type and APP/PS1 mice, both display this intermingled network state, indicating an intact “home cage” network in APP/PS1 mice.

**Figure 2.**
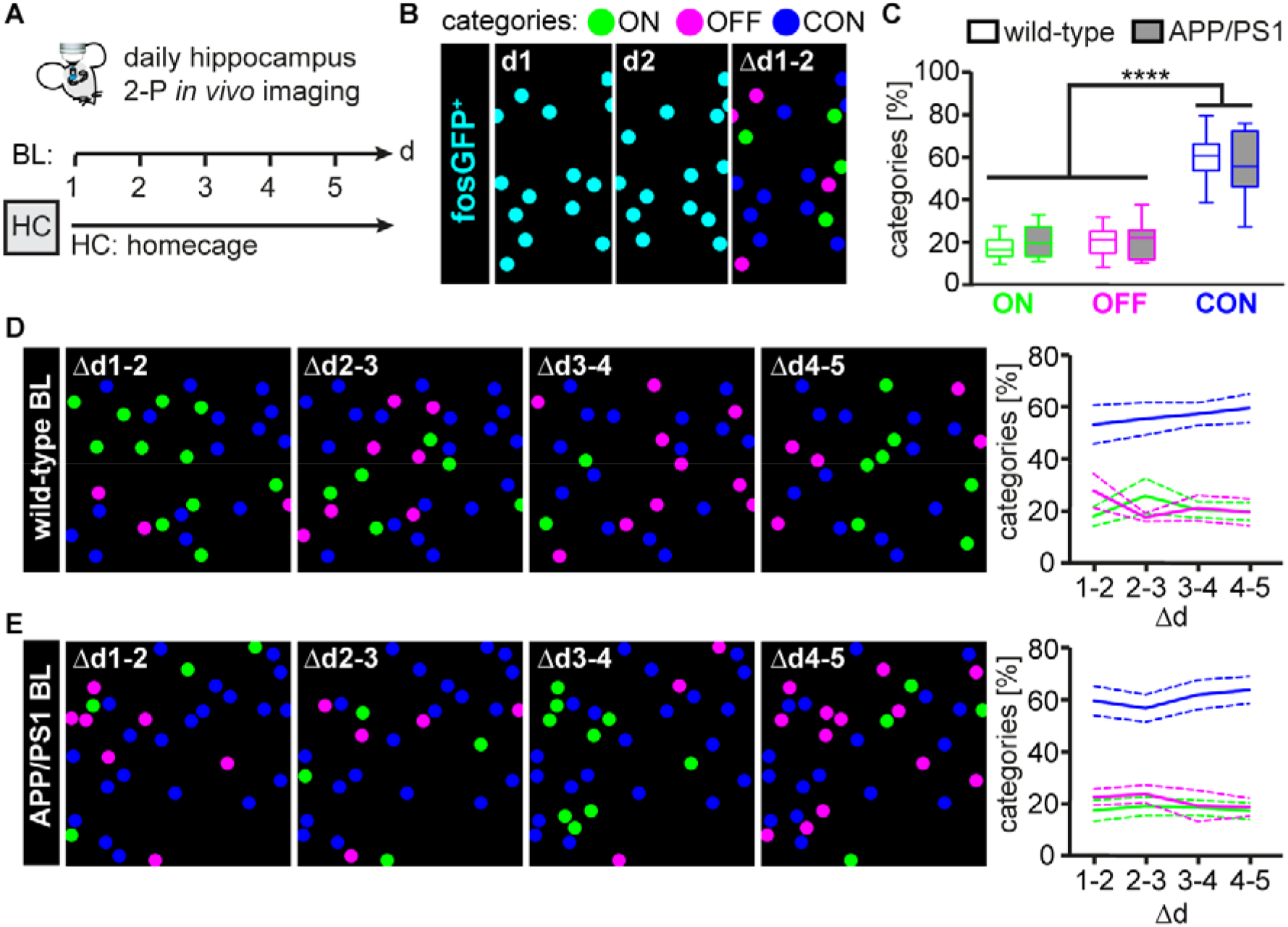
Longitudinal *in vivo* imaging reveals the neuronal network for “home cage” representation. **(A)** Experimental timeline to investigate baseline (BL) fosGFP expression. Images were taken daily, for a period of five days. After each imaging session mice were returned to their home cage (HC). **(B)** Scheme explaining the color-code of daily fosGFP expression changes (d). Neurons that turn on (ON), turn off (OFF) or continue to express fosGFP (CON) were color-coded in green, magenta and blue, respectively. **(C)** Boxplot representing average fractions of ON, OFF and CON neurons over five days in wild-type and APP/PS1 mice. **(D, E)** Representative images illustrating the distribution of ON, OFF and CON neurons and their changes during five days of HC exposure in wild-type (D) and APP/PS1 (E) mice. Right panels: Daily changes of the three categories in wild-type and APP/PS1 mice. Data from n=8 wild-type (4134 fosGFP^+^ neurons) and n=6 APP/PS1 mice (2993 fosGFP^+^ neurons); (C) ****p < 0.0001 (ON vs. CON and OFF vs. CON for wild-type and APP/PS1 mice, respectively), all other comparisons resulted in p-values > 0.05, two-way ANOVA with Holm-Sidak’s correction for multiple comparisons. Data in (D) and (E) are represented as mean (continuous line) ± s.e. m. (dashed lines). See also Figure S2.

### Presence of reactivation pattern independent of memory

We next analyzed the dynamics of CA1 network composition during contextual fear conditioning (cFC), a hippocampus-dependent learning and memory test. We carried out daily *in vivo* imaging of fosGFP^+^ neurons on five consecutive days and performed cFC training and test 90 minutes before the third and fifth imaging time-point, respectively (Fig. 3A). In addition, we analyzed a group of wild-type mice that was trained in context A and tested in a novel context B (Fig. 3A). Both, wild-type and APP/PS1 mice showed similar pre-shock freezing rates and distances traveled, excluding any differences in mobility between genotypes that could bias the results (Supplementary Fig. 3A,B). In accordance with previous results (Kilgore et al., 2010; Roy et al., 2016), APP/PS1 mice trained and tested in context A showed significantly reduced freezing rates compared to wild-type mice (Fig. 3B). Furthermore, wild-type mice trained in context A and tested in a novel context B exhibited significantly decreased freezing rates compared to wild-type mice tested in context A (Fig. 3B). We hypothesized that these differences in freezing behavior are manifested in the number of fosGFP^+^ neurons after memory acquisition in the three different groups. Interestingly, we found an increased number of ON neurons after training in both, wild-type and APP/PS1 mice, suggesting that memory acquisition was intact in the mouse model of AD (Fig. 3C,D,F; Supplementary Fig. 4). This finding was consistent with previous results showing intact memory encoding at early disease stages using the same mouse model (Roy et al., 2016). To exclude any effect of fosGFP expression kinetics, we confirmed that the time period to switch on and off fosGFP expression was reliably within the one-day imaging intervals (Supplementary Fig. 3C,D). On the day between training and test we detected an increased number of OFF neurons in wild-type and APP/PS1 mice, suggesting that the network activity went back to baseline levels (Fig. 3C,D,G; Supplementary Fig. 4). The number of CON neurons remained stable over time in all three experimental groups (Fig. 3CE,H). Retrieval in the conditioned context A did not evoke an increased number of ON neurons in either wild-type or APP/PS1 mice. In contrast, exposing wild-type mice to a novel context B induced another increase of ON neurons (Fig. 3C-F; Supplementary Fig. 4), confirming previous results showing that Fos is mainly triggered by exploration of novel environments, rather than by exploration of familiar environments or aversive stimuli alone (Radulovic et al., 1998). These data suggest an intact fosGFP expression response upon cFC and retrieval comparing wild-type and APP/PS1 mice. However, what else could account for the impaired memory of APP/PS1 mice on the cellular level? To answer that question, we next investigated reactivation of CA1 neurons potentially representing a memory trace and asked whether they were affected in APP/PS1 mice. It has been shown that recent and remote contextual memory retrieval involves the reactivation of partially overlapping subsets of CA1 neurons that have been recruited during learning (Guzowski et al., 1999; Tayler et al., 2013). Optogenetic silencing of excitatory CA1 pyramidal neurons either during acquisition or retrieval prevented memory formation and recall (Goshen et al., 2011). Furthermore, targeted optogenetic stimulation of neuronal ensembles that have been tagged during learning was sufficient to artificially induce memory recall two days after conditioning (Ryan et al., 2015). Based on these findings we hypothesized that APP/PS1 mice exposed to the conditioned context A and wild-type mice tested in the novel context B exhibited decreased reactivation of fosGFP^+^ neurons compared to wild-type mice tested in the conditioned context A. Therefore, we calculated the percentage of reactivated neurons according to the “standard” definition of reactivation: percentage fosGFP^+^ after cFC and test, of fosGFP^+^ after cFC (Fig. 4A). In addition, our data provided information about fosGFP^+^ neurons before cFC and in between cFC and test enabling a “precise” definition of reactivation: fosGFP^+^ after cFC and test, but definitely fosGFP-negative (fosGFP^−^) before cFC and test (Fig. 4B). With the standard definition of reactivation we observed similar fractions of reactivated neurons during the baseline period and the cFC-test period (Fig. 4C). Furthermore, we found no significant difference between the three experimental groups, indicating that the standard definition of reactivation was not sufficient to detect specific memory traces (Fig. 4C). In contrast, using the precise definition of reactivation revealed a significantly increased fraction of reactivated neurons comparing the baseline and cFC-test period (Fig. 4D). Surprisingly, all three experimental groups exhibited similar fractions of these precisely reactivated neurons during learning and memory (Fig. 4D). If the fraction of precisely reactivated neurons in APP/PS1 mice is similar to wild-type mice, what else reflects the memory deficit of APP/PS1 mice on the cellular level? Might the simultaneous activity of other neuronal ensembles interfere with the ensemble of reactivated neurons, thereby preventing memory recall in APP/PS1 and wild-type mice exposed to a novel context? We addressed this question by performing a pattern analysis comparing the frequency of all possible combinations of “active” and “silent” states of individual CA1 neurons during baseline and the cFC-test period for the three experimental groups (Fig. 4E,F; Supplementary Fig. 5). Again, during the cFC-test period all three groups exhibited a comparable frequency of the precise reactivation pattern (Fig. 4F, pattern C), which was elevated compared to the baseline period (Supplementary Fig. 5). Additionally, neurons fosGFP^+^ only after cFC were also increased (Supplementary Fig. 5, pattern L) and again comparable between groups (Fig. 4F). These data indicate that neurons turning on fosGFP expression after training consist of two subsets in all three experimental groups: Firstly, neurons that are active once after training possibly representing novel context information. Secondly, reactivated neurons that are active twice after training and test, presumably representing a memory trace. Interestingly, APP/PS1 mice and wild-type mice tested in context B, the two groups with significantly decreased freezing rates (Fig. 3B), exhibited another increased neuronal subset, composed of new fosGFP^+^ neurons that were specifically detected only after memory testing on day 5 (Fig. 4F, Supplementary Fig. 5, pattern F). The simple comparison of the groups’ relative pattern frequencies during the cFC-test period, revealed a high similarity between APP/PS1 and wild-type mice tested in context B (Fig. 4F). In contrast, the curve of wild-type mice tested in context A exhibited fewer maxima and thereby differed from the two other experimental groups (Fig. 4F; Supplementary Fig. 5). These data suggest the presence of an additional subset of fosGFP^+^ neurons during memory retrieval in the two groups with decreased fear memory as main difference to wild-type mice with intact memory recall.

**Figure 3.**
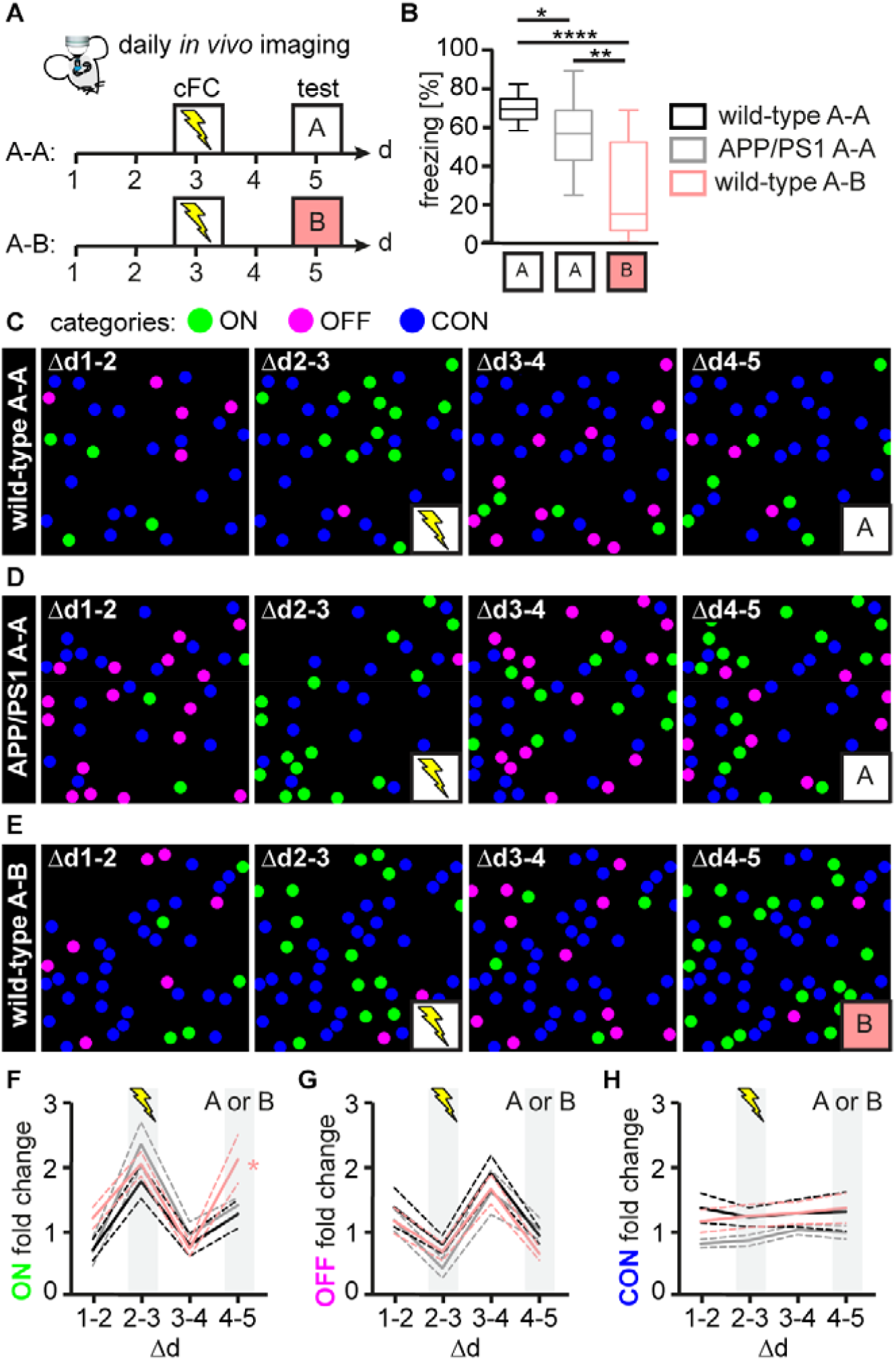
Changes in the composition of fosGFP^+^ CA1 neurons during learning and memory. **(A)** Experimental timeline to investigate the dynamic composition of the hippocampal neuronal network after contextual fear conditioning (cFC) and memory test in the conditioned context A (A-A) or a novel context B (A-B). Training and test were conducted 90 minutes before imaging sessions on days three and five, respectively. **(B)** Average freezing rates of wild-type and APP/PS1 mice during memory test in the conditioned context A or a novel context B. **(C-E)** Representative binary images illustrating the daily changes (d) of ON, OFF and CON neurons of wild-type (C) and APP/PS1 mice (D), trained and tested in context A (AA) and of wild-type mice (E) trained in context A, but tested in context B (A-B). **(F-H)** Intergroup comparisons of daily fold changes of ON (F), OFF (G) and CON (H) neurons during the experimental period. (B) Data from n=16 wild-type A-A, n=14 APP/PS1 A-A and n=10 wild-type A-B mice; *p=0.0322 (wild-type A-A vs. APP/PS1 A-A), **p=0.0014 (APP/PS1 A-A vs. wild-type A-B), ****p<0.0001 (wild-type A-A vs. wild-type A-B), ordinary one-way ANOVA with Holm-Sidak’s correction for multiple comparisons. (F-H) Average values from n=8 wild-type A-A (4775 neurons), n=6 APP/PS1 A-A mice (3776 neurons) and n=6 wild-type A-B mice (5099 neurons); *p=0.0264 (∆d4-5, wild-type A-A vs. wild-type A-B), p=0.0748 (∆d4-5, APP/PS1 A-A vs. wild-type A-B), p=0.6656 (∆d4-5, wild-type A-A vs. APP/PS1 A-A), all other comparisons resulted in p-values > 0.05, two-way ANOVA with Holm-Sidak’s correction for multiple comparisons. See also Figure S3 and Figure S4.

**Figure 4.**
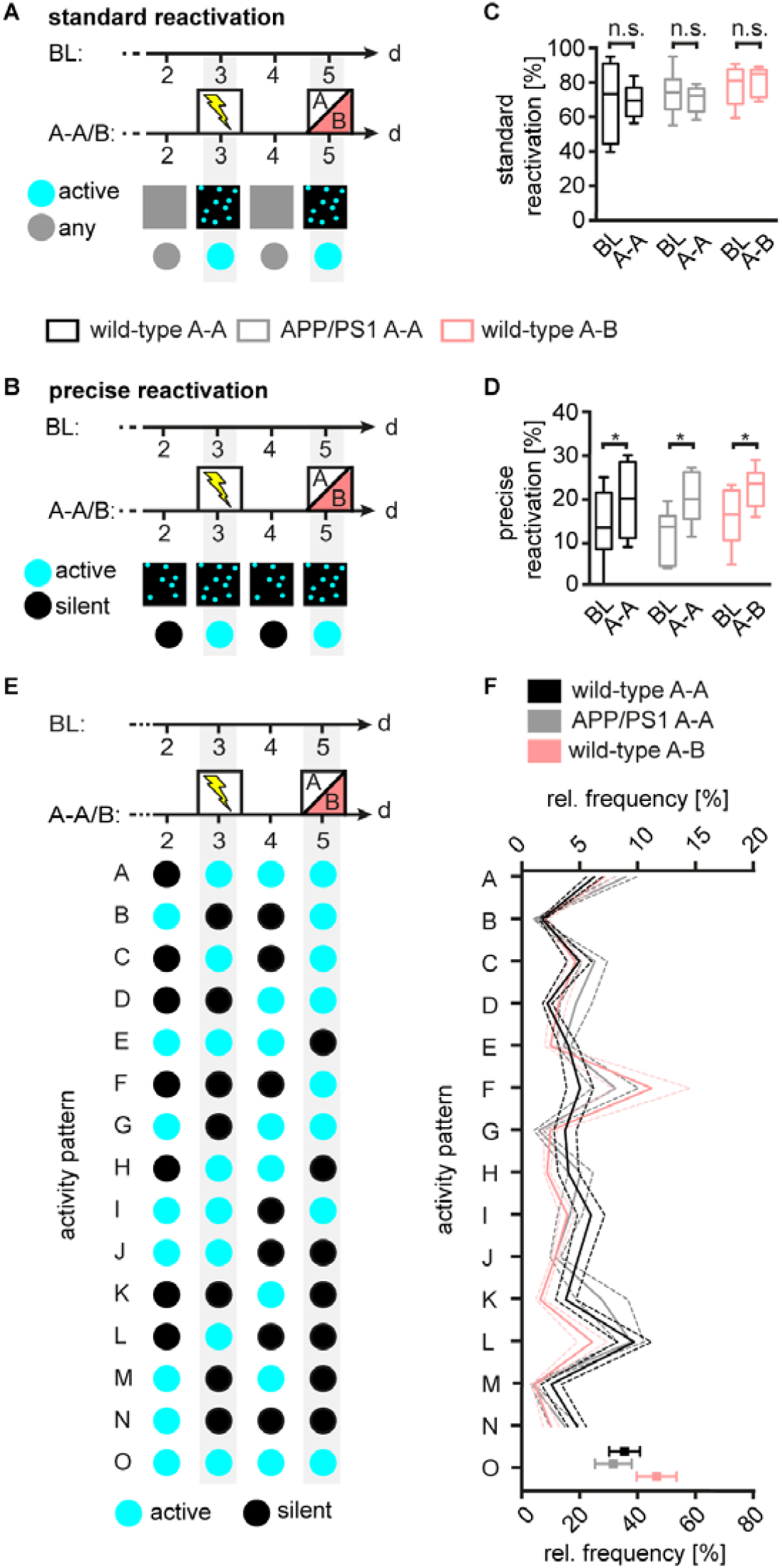
Presence of reactivation pattern independent of memory. **(A, B)** Schematics illustrating the two different definitions of reactivation: (A) “standard reactivation”: neurons express fosGFP (active, cyan) after training and test irrespective of their fosGFP level on the days before (any, grey). (B) “Precise reactivation”: fosGFP expression of neurons switches specifically from fosGFP-negative (silent, black) to active after training and test, respectively. **(C, D)** Intragroup (BL vs. A-A/B) comparison of neuronal subsets with “standard reactivation” (C) or “precise reactivation” (D). **(E)** Illustration of every possible activity pattern an individual neuron can adopt during the baseline (BL) and cFC-test period (A-A/B). **(F)** Relative frequency of every possible activity pattern during the cFC-test period comparing the three experimental groups. Data from n=8 wild-type A-A (4775 neurons), n=6 APP/PS1 A-A mice (3776 neurons) and n=6 wild-type A-B mice (5099 neurons); (C) p-values for intra-group comparisons > 0.05 (n.s.), (D) *p=0.0341 (wild-type A-A), *p=0.0127 (APP/PS1 A-A), *p=0.0368 (wild-type A-B), two-way ANOVA with correction for multiple comparisons using the two-stage linear step-up procedure of Benjamini, Krieger and Yekutieli. See also Figure S5.

### Impurity of the memory trace decreased memory retrieval

So far we demonstrated the existence of reactivated fosGFP^+^ neurons in all three experimental groups independent of memory recall performance, indicating intact memory acquisition in the mouse model of AD. Therefore, we hypothesized that memory retrieval impairment manifested in another neuronal subset, different from reactivated neurons. To test this hypothesis, we analyzed the activity history of neurons that were fosGFP^+^ after retrieval, constituting the retrieval network (RN, Fig. 5A). As expected, the RN contained an increased fraction of fosGFP^+^ neurons with reactivation history (REAH) in all three groups (Fig. 5A,B,D). However, another activity history pattern with the highest frequency increase found in APP/PS1 tested in the conditioned context A and wild-type mice tested in the novel context B comprised neurons only activated after retrieval (RONLY pattern, Fig. 5A,C,D). In contrast, wild-type mice tested in context A exhibited a frequency decrease of the RONLY pattern (Fig. 5A,C,D). These data indicate that APP/PS1 mice tested in context A and wild-type mice tested in context B possessed two major elevated neuronal populations in their retrieval network: First, neurons with the REAH pattern and second, neurons with the RONLY pattern. This superimposition of the potential memory trace REAH by the RONLY pattern presumably reflects the decreased memory on the neuronal network level. To compare the entire RN composition of the three experimental groups, we included all possible activity history patterns in our analysis (Fig. 5A). We generated a network similarity map by sorting the patterns according to their frequency change in descending order and connected corresponding patterns of each experimental group with a line. The more intersections appeared, the higher the dissimilarity of the compared retrieval network. The less intersections occurred, the higher the similarity (Fig. 5E). There was a striking difference between the RN of wild-type mice with intact memory to the other groups (Fig. 5E). Surprisingly, the RNs of APP/PS1 mice tested in context A and wild-type mice tested in context B were very similar to each other indicating that APP/PS1 mice perceive the re-exposure to the conditioned context A as an exposure to a novel context. In spite of the presence of a memory trace, APP/PS1 mice tested in context A and wild-type mice tested in context B do not recall the context properly. This was reflected on the RN level, where the memory trace REAH was superimposed by the RONLY trace, which presumably contributed to and/or reflected memory impairment in APP/PS1 mice (Fig. 5F). To test this hypothesis of a superimposed memory trace, we artificially induced CA1 neuronal activity during memory retrieval (Fig. 6). We targeted the expression of an activating DREADD (designer receptor exclusively activated by designer drug; hM3Dq) to dorsal CA1 by AAV-mediated delivery of cre/*loxP-*dependent DREADD expression in Vglut2-ires-cre mice restricting expression to excitatory glutamatergic neurons (Fig. 6A). Two days after contextual fear conditioning mice were tested in the conditioned context with previous clozapine-N-oxide (CNO) or placebo injections, respectively. 90 minutes after memory retrieval mice were perfused to immunohistochemically stain for endogenous Fos expression. Confocal images of coronal brain sections revealed specific hM3Dq expression in the hippocampal CA1 region (Fig. 6C,D). CNO induced hM3Dq-mediated activation of CA1 neurons as indicated by a significant increase of Fos^+^ nuclei within CA1, compared to placebo-injected control mice (Fig. 6C-E). Mice with artificially induced neuronal activity exhibited decreased memory retrieval performance, indicated by a decreased freezing rate (Fig. 6F). These data show that artificial non-specific activation of excitatory neurons in the CA1 neuronal network is sufficient to decrease recall performance. We next asked whether artificially induced false context information alone would cause impaired retrieval in wild-type mice (Fig. 7). To test this, we utilized the TetTag-system (Zhang et al., 2015) to express hM3Dq after induction of the c-fos promoter in an activity-dependent manner (Supplementary Fig. 6). The AAV combination (AAV-Fos-tTA and AAV-PTRE-tight-hM3Dq-mCherry) was injected bilaterally into CA1 (Fig. 7A). Doxycycline (DOX) was present in the drinking water to prevent unspecific expression of hM3Dq. Neurons in CA1 encoding the information of context B (false context) were labeled with hM3Dq-mCherry by setting mice off DOX (Fig. 7B) and exposing them to context B. Control mice were kept on DOX during context B exposure (non-labeled). Two days after contextual fear conditioning mice were tested in the conditioned context with previous clozapine-N-oxide (CNO) or placebo injections, respectively. Fos staining revealed an increased CNO-mediated activation of hM3Dq-expressing CA1 neurons as indicated by a significant increase of Fos^+^ nuclei within CA1, compared to placebo-injected and non-labeled control mice (Fig. 7C-E, F). Mice with artificially superimposed false context information during memory retrieval exhibited decreased freezing rates (Fig. 6G). These data suggest that superimposition of false context information during retrieval of the conditioned context A is sufficient to decrease memory.

**Figure 5.**
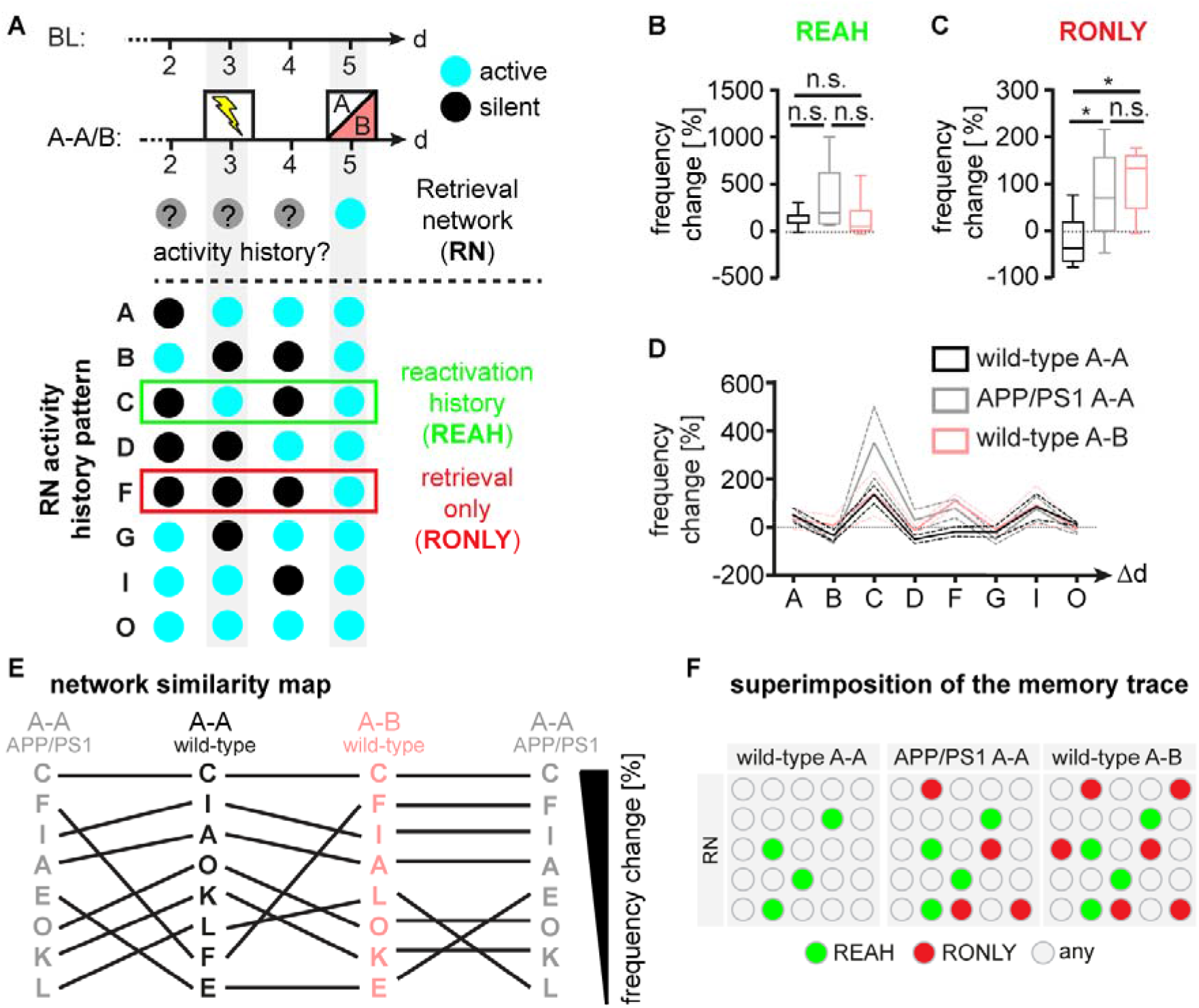
Aberrant retrieval network (RN) composition in APP/PS1 mice. **(A)** Scheme illustrating the activity history analysis of fosGFP^+^ (active) neurons constituting the RN. Illustrated are neurons with a reactivation history (REAH, green box) and neurons without past fosGFP expression, which are fosGFP^+^ after exposure to the retrieval context only (RONLY, red box). **(B)** Change in frequency of the REAH pattern between BL and cFC-test periods (A-A/B) for the three experimental groups. **(C)** Same as (B) for RONLY neurons. **(D)** All possible RN activity history pattern including REAH (pattern C) and RONLY (pattern F) pattern. **(E)** Network similarity map: frequency change of each pattern between BL and cFC-test period (A-A/B) in descending order. Corresponding pattern of the experimental groups are connected with lines. The higher the similarity the fewer the intersections of connecting lines. **(F)** Model illustrating the superimposition of the memory trace in APP/PS1 mice. Data from n=8 wild-type A-A mice (3782 cells), n=6 APP/PS1 A-A mice (2538 cells), and n=6 wild-type A-B mice (4105 cells); (B) inter-group comparisons p>0.05 (n.s.), one-way ANOVA with Holm-Sidak’s correction for multiple comparisons; (C) *p=0.0426 (wild-type A-A vs. APP/PS1 A-A), *p=0.0109 (wild-type A-A vs. wild-type A-B), p=0.4450 (APP/PS1 A-A vs. wild-type AB), ordinary one-way ANOVA with Holm-Sidak’s correction for multiple comparisons.

**Figure 6.**
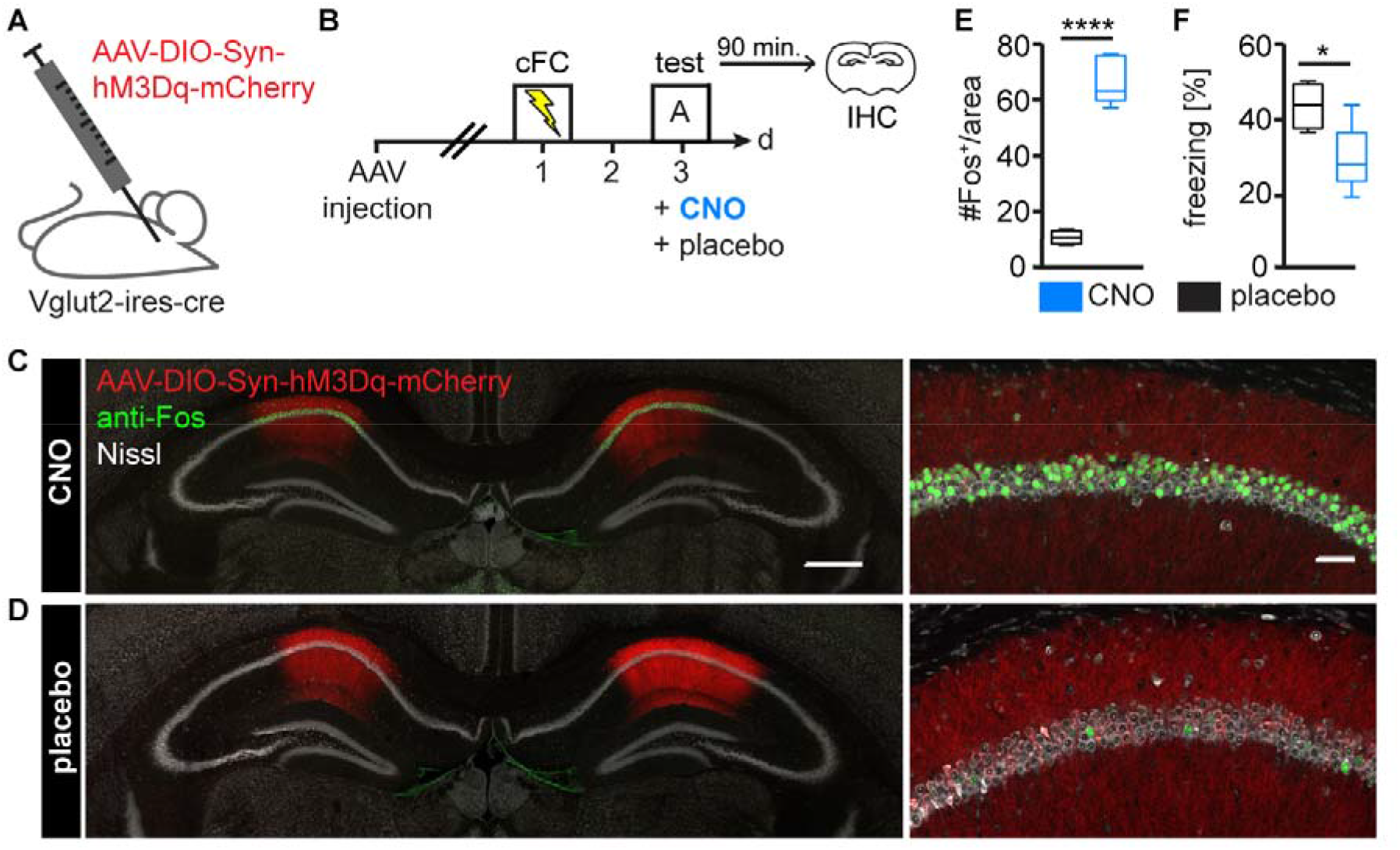
Disturbing the RN by superimposition of the memory trace decreases recall performance. **(A, B)** Schematic of hM3Dq-mediated activation during memory retrieval to superimpose the memory trace. The activating DREADD hM3Dq was injected into CA1 bilaterally, targeted to excitatory glutamatergic neurons via Cre/*loxP*-dependent expression in Vglut2-ires-cre mice (A). Experimental timeline for accessing memory retrieval performance during superimposition of the memory trace (B). After three weeks of recovery from AAV injection, mice were fear conditioned on d1 and tested in the conditioned context on d3 with an injection of clozapine-N-oxide (CNO) or placebo, respectively, 40 minutes prior to memory test. **(C, D)** Typical confocal overview (left) and detailed (right) images of coronal sections showing bilateral expression of hM3Dq-mCherry and Fos in CA1 of mice injected with CNO
(C) or placebo (D), prior to memory test. **(E)** Comparison of Fos^+^ nuclei within CA1 of the CNO- and placebo-treated groups. Nuclei within an area of 300 µm x 80 µm were counted. **(F)** Freezing of CNO- and placebo-treated mice during memory test. Data from n=5 CNO-treated and n=4 placebo-treated mice. (E) ****p<0.0001, unpaired t-test; (F) *p=0.0112 unpaired t-test with Welch’s correction. Scale bars: (C) 500 µm (left), 50 µm (right).

**Figure 7.**
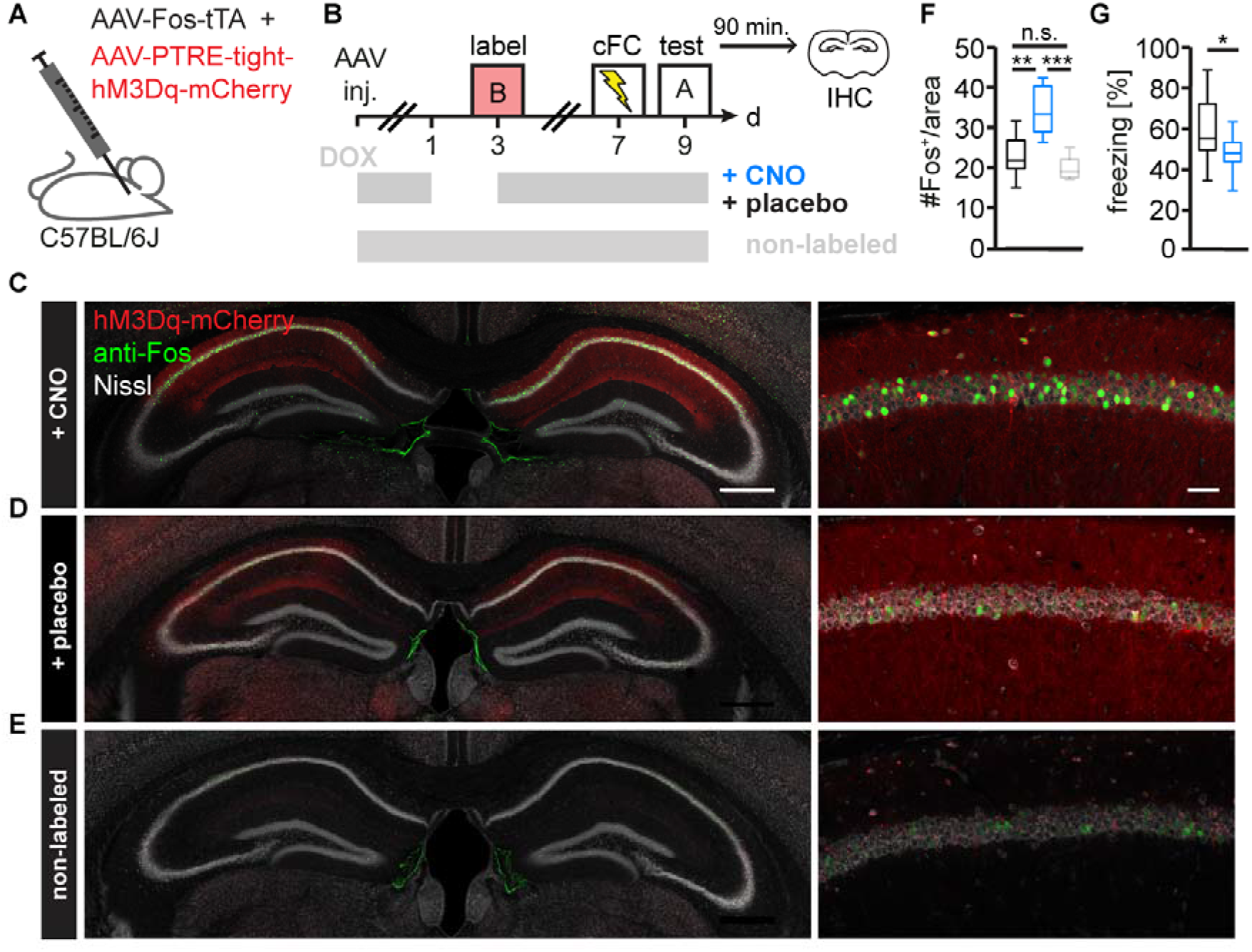
Memory superimposition by false context information impairs memory. **(A, B)** Schematic of memory trace superimposition by false context information via activity-dependent hM3Dq expression. AAVs containing the TetTag system were injected into CA1 bilaterally (A). Experimental timeline for accessing memory retrieval performance during superimposition of the memory trace (B). Mice were kept on doxycycline (DOX) from the day of AAV injections on. After three weeks of recovery, mice were depleted from DOX for two days for labeling neurons encoding the novel context B (false context) on d3. Control mice were also exposed to the false context, but with continuous DOX garbage (non-labeled). All mice were fear conditioned on d7 and tested in the conditioned context on d9 with an injection of clozapine-N-oxide (CNO) or placebo, respectively, 40 minutes prior to memory test. IHC, immunohistochemistry. **(C, D, E)** Exemplary confocal overview (left) and detailed (right) images of coronal sections showing bilateral expression of hM3Dq-mCherry and Fos in CA1 of mice injected with CNO (C) or placebo (D) prior to memory test, and non-labeled mice (E). **(F)** Comparison of Fos^+^ nuclei within CA1 of the CNO- and placebo-treated groups, and non-labeled mice. Nuclei within an area of 300 µm x 80 µm were counted. **(G)** Freezing of CNO- and placebo-treated mice during memory test on d9. Data from n=12 CNO-treated and n=12 placebo-treated mice. (F) **p=0.0018 (placebo vs. CNO), ***p=0.0005 (CNO vs. non-labeled), p=0.2643 (placebo vs. non-labeled), ordinary one-way ANOVA with Holm-Sidak’s correction for multiple comparisons; (G) *p=0.0379 unpaired t-test. Scale bars: (C) 500 µm (left), 50 µm (right). See also Figure S6.

## Discussion

We monitored fosGFP expression dynamics in the hippocampal CA1 region *in vivo* and investigated memory trace encoding neurons during a contextual fear conditioning test in wild-type mice and a mouse model of AD. We demonstrated that the CA1 network contains two main subsets of neurons – one continuously active and one recruited and released from the network on a daily basis. Upon memory acquisition we detected a substantial amount of neurons active for the first time. Only a fraction (20%) of these neurons were specifically reactivated after memory retrieval, indicating that the memory trace consisted of a smaller but specific subset of CA1 neurons. Surprisingly, we identified memory traces in both wild-type mice and in a mouse model of AD with impaired memory. This result posed the question, what hampered memory retrieval despite the existence of reactivated neurons in the AD mouse model? By performing an activity pattern and network similarity analysis, we revealed a subset of neurons only active after retrieval of the conditioned context in APP/PS1 and wild-type mice exposed to a novel context. Indeed, the retrieval network composition of APP/PS1 mice resembled that of wild-type mice tested in a novel context, which was characterized by neurons newly recruited to the retrieval network, superimposing the memory trace. Indeed, mimicking memory trace superimposition by artificially activating false context information in CA1 during retrieval decreased memory recall performance. These data support the conclusion that memory retrieval impairment in the AD mouse model is reflected on the neuronal network level by superimposition of an intact memory trace.

IEG expression in neurons has been used as a correlate of recent neuronal activity and hence, to identify memory traces (Gall, 1998; Guzowski et al., 1999; Josselyn et al., 2015; Rogerson et al., 2014; Silva et al., 2009). In the current study, we used a mouse model that expressed GFP under the control of the promoter of the IEG *c-fos (Barth et al., 2004)*. The expression of fosGFP in this mouse model was shown to reliably report neuronal activity in the cortex *in vivo* (Jouhanneau et al., 2014; Yassin et al., 2010) and the hippocampus *in vitro* (Schoenenberger et al., 2009). In line with these results, we found increased fosGFP expression 1.5 h after contextual fear conditioning which mirrored endogenous Fos protein levels. In AD and mouse models thereof epileptiform neuronal activity has been detected (Palop et al., 2007; Vossel et al., 2013). This was supported by the finding of hyperactive neurons in the cortex and hippocampus of AD mouse models (Busche et al., 2012; Busche et al., 2008; Rudinskiy et al., 2012). Correspondingly, using the IEG *arc* as reporter for neuronal activity in the visual cortex revealed a subset of neurons with increased *arc* expression in proximity to A plaques (Rudinskiy et al., 2012). In contrast, we found a decrease of fosGFP expression in proximity to A plaques indicating decreased neuronal activity dependent on A deposition. Whether these discrepancies are attributable to the different mouse models, brain regions or methods to determine neuronal activity (Ca^2+-^imaging, *arc*, *c-fos*) remains to be further characterized. Indeed, the same study that found hyperactive neurons also revealed a large fraction of hypoactive hippocampal neurons in the AD mouse model (Busche et al., 2012). In summary, both hypo- and hyperactivity argue for an imbalanced hippocampal network that forms the basis for memory deficits under AD-like conditions.

To understand the dynamic composition of the hippocampal neuronal network beyond a classical two time-points analysis, we utilized longitudinal two-photon *in vivo* imaging of individual fosGFP^+^ neurons in mice. A two time-points analysis of *cfos*-driven reporter expression was used to detect reactivated neurons and to identify memory traces (Reijmers et al., 2007; Tayler et al., 2013). Beyond that, longitudinal *in vivo* imaging enables reporter expression measurements of individual neurons at many time-points and with spatial relationship (Trachtenberg et al., 2002). This allows studying the changes of a neuronal network composition with superior spatio-temporal resolution. Using this technique, we revealed two sub-populations of hippocampal CA1 neurons: one constantly expressing fosGFP (60%) and one turning fosGFP expression on and off on a daily basis (20% each). This result underscores on the one hand the plasticity of the neuronal network, but on the other hand emphasizes the existence of a backbone or default network on top of which new information is processed. These data support the view that newly formed memories have to be integrated in a network composed of preexisting information (Josselyn et al., 2015; Rogerson et al., 2014). The default network was intact in the AD mouse model indicating that other alterations underlie learning and memory deficits. Therefore, we next analyzed changes of the neuronal network activity in hippocampal CA1 area during contextual fear conditioning and retrieval. In accordance with previous studies in wild-type and APP transgenic mice, we revealed an increased number of fosGFP^+^ neurons in hippocampal CA1 area after contextual fear conditioning (Palop et al., 2005; Radulovic et al., 1998). These data pointed to a defective memory retrieval rather than memory acquisition in APP/PS1 transgenic mice. Indeed, our activity pattern analysis revealed that memory traces that consisted of reactivated neurons were present in both, mice with memory (wild-type A-A) and mice with decreased memory (APP/PS1 A-A and wild-type A-B), underscoring the integrity of memory acquisition. Of note, only a fraction of neurons (20%) that turned on fosGFP after cFC were reactivated after retrieval indicating that the actual memory trace comprised a much smaller but refined subset of neurons than initially activated during acquisition. This finding is intriguing against the background of recent theories about memory consolidation – obviously, memory traces in the hippocampal CA1 area are rapidly refined presumably increasing the efficiency of information processing (Josselyn et al., 2015).

Why were memory traces present independent of memory? By integrating the past activity history we were able to identify an additional subset of fosGFP^+^ neurons that was only present during retrieval in the two groups with decreased memory. This subset of neurons presumably superimposed the memory trace and interfered with effective recall. To artificially induce memory trace superimposition, we induced unspecific neuronal activity during memory retrieval by pharmacogenetically activating excitatory glutamatergic neurons in hippocampal CA1. Indeed, this artificially induced activity reduced memory retrieval performance underscoring the importance of the presence of a pure memory trace for effective recall. Furthermore, superimposing the memory trace by artificially activating CA1 neurons coding a specific other context lead to similar impaired memory as unspecific excessive activation. These findings are in line with previous experiments demonstrating that brain-wide artificial pharmacogenetic activation of neurons encoding a novel context during retrieval in the conditioned context decreased memory retrieval performance (Garner et al., 2012). Our data extent this finding by revealing that activation of a false memory during recall restricted to the CA1 area is sufficient to induce memory impairment.

The hippocampal CA1 area has been proposed to play an important role in detecting novelty due to its ability to act as a comparator between past and present experiences (Kumaran and Maguire, 2007). Our data suggest a defective comparator in APP/PS1 mice leading to a false mismatch detection and thus, novelty-induced CA1 activity during retrieval in an actually familiar environment. It has been shown that entorhinal rather than CA3 inputs are the main drive for this novelty-induced CA1 activity (Karlsson and Frank, 2008). Furthermore, the medial entorhinal cortex (EC) is affected early during pathology of Alzheimer’s disease (Braak and Braak, 1991; Hsia et al., 1999) and optogenetic stimulation of connections between dentate gyrus (DG) and EC was sufficient to rescue memory deficits in the same mouse model of AD (Roy et al., 2016). Together with our data, this points at impaired or altered EC-CA1 connectivity in APP/PS1 mice. In summary, we demonstrate that memory traces consist of only a small subset of reactivated neurons. Our data support that memory acquisition of APP/PS1 transgenic mice is intact, which is reflected on the neuronal network level by the formation and presence of memory traces in CA1. Memory retrieval is affected due to superimposition of the memory trace by aberrantly active neurons during memory recall. Artificial induction of false context information in CA1 of healthy mice during retrieval was sufficient to mimic the memory deficit in APP/PS1 mice. These results will be helpful for the future development of interventions directed at restoring the purity of memory traces in human AD patients.

## Author contributions

S.P. conducted the experiments, analyzed the data, prepared figures and wrote the manuscript. L.S., J.S., J.W. provided technical assistance. D.E. and W.S.J. provided animal models and conceptual advice on experiments and manuscript. S.S. produced and provided AAVs. B.S. synthesized and provided reagents. M.F. wrote the manuscript, coordinated research and supervised the project.

## Acknowledgements

This work was supported by the DZNE, grants from the Deutsche Forschungsgemeinschaft (SFB 1089, KFO 177), Centres of excellence in Neurodegeneration (CoEN). We thank P. Thevenaz and Erik Meijering for the development of the ImageJ plugins “stackreg” and “TurboReg”. We thank Simon Wiegert, Stefan Remy and Gabor Petzold for helpful discussion on the manuscript and the light microscopy facility of DZNE for constant support.

## Methods

### Transgenic mice

FosGFP and APP_swe_/PSEN1dE9 (APP/PS1) mice were obtained from The Jackson Laboratory (Stock number: 014135) and the Mutant Mouse Regional Resource Center (Stock number: 034832; formerly JAX Stock No. 005864) respectively. Both mouse lines were maintained on a C57/BL6 background. FosGFP mice express enhanced green fluorescent protein (EGFP) under the *c-fos* promoter resulting in a fosGFP fusion protein. APP/PS1 transgenic mice express human amyloid precursor protein with a Swedish mutation APPK595N/M695L under the mouse prion protein promoter and a mutant human presenilin1 with a deletion of exon 9 (PS1-dE). Heterozygous FosGFP^tg/wt^ and APP/PS1^tg/wt^ were crossbred. Experiments were carried out at an age of 13-16 months. *Slc17a6^tm2(cre)Lowl^*/J (VGlut2-ires-cre) mice (The Jacksons Laboratory, Stock number: 016963) were on a 129S4 background and 6-8 months of age. Mice were group-housed in colonies of up to 5 mice, separated by gender in IVC cages under specific pathogen free (SPF) conditions with unlimited access to food and water. The light and dark cycle was 12h/12h and the temperature was kept constant at 22°C. Equal numbers of male and female mice were randomly assigned to the experimental groups. Behavioral experiments were carried out at the dark cycle, imaging was carried out at the light cycle, respectively. All procedures were in accordance with an animal protocol approved by the DZNE and the government of North-Rhine-Westphalia.

### Hippocampal window surgery

We surgically implanted a unilateral cranial window on top of the dorsal hippocampus as previously described (Gu et al., 2014; Schmid et al., 2016). The mice were anesthetized with an intraperitoneal injection of ketamine/xylazine (0.13/0.01 mg/g body weight). Additionally, an anti-inflammatory (dexamethasone, 0.2 mg/kg) and an analgesic drug (buprenorphine 0.05 mg/kg; TEMGESIC^®^, Reckitt Benckiser Healthcare (UK) Ltd., Great Britain) were subcutaneously administered immediately before surgery. The analgesic was applied for three consecutive days after surgery as post-operative treatment. To avoid an experimental bias caused by inflammatory processes, mice were given at least three weeks to completely recover from surgery before *in vivo* imaging started (Gu et al., 2014).

### AAV injection

For bilaterally injecting AAVs, mice were anesthetized, the hair was removed at the site of incision and the skin was wiped with 70% ethanol. A small incision was made at the midline to expose bregma and the sites of injection. Small holes were drilled and the opening of the dura was ascertained. From here the needle was lowered to the depth of interest from brain surface. A volume of 0.5 µL virus per hemisphere was injected with a speed of 0.1 µL/minute. After injecting the virus the needle was left at the site of injection for 10 minutes to allow the virus to diffuse into the tissue. The skin was closed by stitches and mice were given three weeks to recover before experiments started. Coordinates for targeting CA1 with AAV-DIO-hM3Dq-mCherry (UNC Vector Core, titer: 6.1*10^12^ virus genomes/mL) were −1.85 mm (AP), ± 1.50 mm (ML), −1.10 mm (DV). For targeting CA1 with the combination of AAV-Fos-tTA and AAV-PTRE-tight-hM3Dq-mCherry (1:2, produced and provided by Susanne Schoch) were −1.95 mm (AP), ± 1.50 mm (ML), −1.15 mm (DV).

### Hippocampus two-photon *in vivo* imaging

*c-fos*-driven EGFP expression data were acquired using an upright Zeiss (Carl Zeiss Microscopy GmbH, Jena, Germany) Axio Examiner LSM7MP setup, equipped with a Coherent Cameleon Ultra2 two-photon laser (Coherent, Dieburg, Germany). A 16x water immersion objective with a numerical aperture of 0.8 (Nikon) was applied. Enhanced green fluorescent protein (EGFP) was excited at 920 nm. Fluorescence emission was separated by a dichroic mirror (LP555), detecting the green (BP 500-550) and red emitted light (BP 575-610) with non-descanned detectors (NDDs). Image acquisition was performed with ZEN2010 (Carl Zeiss Microscopy GmbH). For each imaging session mice were anesthetized with ketamine/xylazine (0.13/0.01 mg/g body weight). To record fosGFP expression in the dorsal CA1 region of the hippocampus we acquired a tile scan consisting of 3 x 3 separate z-stacks of 120 µm depth with 3 µm z-resolution, starting at the surface of the stratum pyramidale. This resulted in an total scan area of 4.57 mm^2^ (2.013 pixels/µm) with a depth of 0.12 mm (total volume of 0.55 mm^3^) spanning a large part of the dorsal CA1 area. For each mouse one to three regions of interest (500 µm x 500 µm) were analyzed. Repetitive scanning of the same positions over time was achieved by orienting to the vascular pattern under reflected light illumination using a GFP filter set and a metal halide lamp HXP100 (Carl Zeiss Microscopy GmbH, Jena, Germany). Visualization of MeXO4-stained A plaques was achieved by exciting at 780 nm and detecting emission with a BP 450/60 filter. Contextual fear conditioning and memory test were conducted 1.5 hours before imaging sessions to ensure induction of fosGFP expression. To avoid handling-evoked fosGFP expression mice were accustomed to the experimenter and to the involved transport between holding and behavior rooms on three consecutive days before the first imaging session. For *in vivo* imaging mice were anaesthetized and let recover from anesthesia in their home cage. APP/PS1 transgenic mice received a dose of 50 µg MeXO4 (0,5 µg/µL MeXO4, 10% DMSO, 45% 1,2-Propanediol) (Burgold et al., 2011) during the handling sessions. Wild-type mice received the same volume of a placebo (solvent without MeXO4).

### Analysis of two-photon *in vivo* images

Raw data z-stacks were processed with the open source software Fiji (ImageJ 1.46j). Autofluorescence was reduced by subtracting the red from the green channel. Z-stacks spanning the dorsal hippocampal CA1 area (30 slices with 3 µm z-spacing) were maximum intensity projected. Maximum intensity projections (further referred to as images) of every imaging session were aligned in x-y-dimension using the “TurboReg” plugin (Thevenaz et al., 1998). Fluorescence intensities of the background (BG) and of fosGFP^+^ nuclei were measured within circular masks (Ø 7.45 µm) that were manually placed besides and above fosGFP^+^ nuclei, respectively. Masks were consecutively numbered, thereby enabling to track fosGFP fluorescence intensity of individual neurons throughout the experiment. To categorize neurons into fosGFP^+^ (active) and fosGFP^-^ (silent) a threshold (TH) was set, determined by the mean BG fluorescence plus six times the standard deviation. Obtained binary images were used to calculate the density of fosGFP^+^ neurons. Density and fosGFP fluorescence intensities were measured on day one (d1) of the experimental timeline. To distinguish between fosGFP^+^ neurons in proximity and distant to a MeXO4-stained Aβ plaques, a circular mask with a diameter of 50 µm was centered around the Aβ-plaque. Neurons inside the mask were defined as proximal (<50 µm, near) to the Aβ-plaque, neurons outside as distant (>50 µm, far).

### Visualization of fosGFP expression changes

FosGFP expression changes reflect alterations in neuronal activity in a one-day measurement interval. Changes were categorized in ON (from silent to active), OFF (from active to silent) and CON (staying active). The fractions of ON, OFF and CON neurons referred to the sum of all neurons per day. To calculate the normalized fold change of neuronal categories each obtained value was divided by the corresponding average baseline value. Images containing ON, OFF and CON neurons were generated by pseudo-coloring the circular masks with green, magenta and blue, respectively.

### Activity pattern analysis

For activity pattern analysis we determined the fosGFP expression on four consecutive days from day two to five during the cFC-test period. Each neuron can theoretically adopt 16 different activity pattern (2^4^=16; Fig. 4E). We determined the activity pattern for each neuron and calculated the relative frequency of each pattern in relation to the total number of fosGFP^+^ neurons.

### Network similarity analysis

To compare the retrieval network (RN) of the experimental groups, the frequency of any activity history pattern among the neurons composing the retrieval network (fosGFP^+^ neurons on day 5 in A-A/B period) was calculated. The change in frequency was obtained by referring to the corresponding frequency during baseline. Subsequently, we sorted the pattern in descending order according to their change in frequency. Corresponding patterns of the three experimental groups were connected by lines resulting in a network similarity plot. The fewer the intersections between two compared groups, the more similar were their network adaptations that occurred in the cFC-test period (A-A/B).

### Contextual fear conditioning

The training and the retrieval for contextual fear conditioning were conducted in a chamber (21,5 x 20 x 25 cm) composed of transparent plastic walls and a stainless steel grid floor connected to an aversive stimulator/scrambler (Med associates Inc.). For training mice were placed in the chamber for 120 seconds before the first foot shock was delivered (0,75 mA, 2 seconds). With an inter-shock period of 60 seconds two more shocks were applied and mice were returned to their home cage 60 seconds after the third shock. For contextual memory retrieval mice were placed in the conditioning chamber 48 hours after the fear conditioning and were allowed to explore the context for 5 minutes. After every trial the chamber was cleaned by 70% ethanol. The novel context B was located in a different room than the conditioning chamber. It was composed of red transparent plastic walls and a white soft plastic floor. Here, inter-trial cleaning was conducted with pine flavored 70% ethanol.

During training and retrieval mice were video recorded from above. As readout the travelled distance was analyzed automatically with EthoVision XT (Noldus). In addition, an experimenter blind to the experimental groups determined the cumulative duration of freezing behavior during the first four minutes of the retrieval session, manually. Freezing was defined as the complete absence of movement, except for breathing.

### Tagging neuronal ensembles

Doxycycline (DOX) was delivered in the drinking water at 2 mg/mL and 5% sucrose from the day of injection on (Zhu et al., 2007). For tagging, mice received usual tab water without DOX for two days before they were exposed to the neutral context B for ten minutes. DOX treatment was continued immediately after context B exposure. To control for DOX efficiency in preventing the expression of hM3Dq, a group of mice received DOX, continuously (non-labeled).

### DREADD activation

For activating hM3Dq, 3 µg/g bodyweight CNO (0.4 µg/µL CNO in 0.9% saline, 1% DMSO) was injected intraperitoneally, 40 minutes before start of the memory test. Control animals received a placebo containing just the solvent.

### Immunohistochemistry

Mice were transcardially perfused with phosphate buffered saline (PBS) pH7.4 followed by 4% paraformaldehyde (PFA) for 5 minutes. Brains were fixed over night in 4% PFA in PBS. Slices of 100 µm thickness were cut on a vibratome (Leica VT 1200, Leica Germany). Free-floating slices were permeabilized over night in 0.8% Triton-X100™ and blocked with 4% normal goat serum (ThermoFisher Scientific) plus 4% bovine serum albumin (Carl Roth). Endogenous Fos protein was labeled with an antibody against Fos (1:500; sc-52, Santa Cruz Biotechnology). Inhibitory interneurons were stained with a primary antibody targeting GAD67 (1:1000; MAB5406, Merck Millipore). Secondary labeling was performed with an antibody Alexa Fluor^®^ 488 anti-rabbit and Alexa Fluor^®^ 647 anti-mouse, respectively (both 1:400; ThermoFisher Scientific).

### Confocal microscopy

Confocal images were acquired using an inverted LSM700 microscope (Carl Zeiss Microscopy GmbH, Jena, Germany) with a 20x air objective. Alexa Fluor^®^ 488 and fosGFP were exited at 488 nm and detected using a 490-555 nm bandpass filter. Alexa Fluor^®^ 647 was excited at 639 nm and detected with a long pass 640 nm filter. MeXO4 was excited at 405 nm and detected at a wavelength of up to 490 nm. The pinhole was set to an airy unit of one. Stacks were acquired with a resolution of 0.31 µm/pixel and a z-spacing of 1 µm.

### Statistics

Statistical analysis and preparation of graphs was performed in GraphPad Prism7 (GraphPad Software Inc., La Jolla, USA). We did not apply statistical methods to predetermine sample sizes, but our chosen sample sizes are similar to those generally employed in the field. Subjects were assigned to experimental groups before data acquisition. Assignment was determined by genotype and by aiming at a balanced proportion of sexes in each group. For data collection blinding was not possible. Data analysis was performed blind to the conditions of the experiment via encrypting file names. Mice were excluded from analysis if acquired data sets were incomplete. All box plots report the median, 25%- and 75%-quartile, with whiskers depicting minimum and maximum values of the data. Pattern frequencies and data over time are represented by line diagrams reporting the mean ± standard error of the mean (s.e.m.), by continuous and dashed lines, respectively.

All data sets were tested for normality with the D’Agostino and Pearson normality test (if n>6) or Shapiro-Wilk normality test (if n<6). Intra-group comparisons of normally distributed data were made using a two-way ANOVA with Holm-Sidak’s correction for multiple comparisons. Inter-group comparisons of normally distributed data were made by applying a one-way ANOVA with correction for multiple comparisons using Holm-Sidak’s or the two-stage step-up method of Benjamini, Krieger and Yekutieli. Comparison of two not normally distributed data sets was conducted with the Mann-Whitney test; for comparing more than two not normally distributed data sets the Kruskal-Wallis test with Dunn’s correction for multiple comparisons was applied. All statistical test applied in this study were two-sided. Figures were prepared with Illustrator CS5 Version 15.0.1 (Adobe).

### Data availability

Datasets are available from the corresponding author upon request.

## Supplemental Data

**Figure S1. Related to Figure 1.**
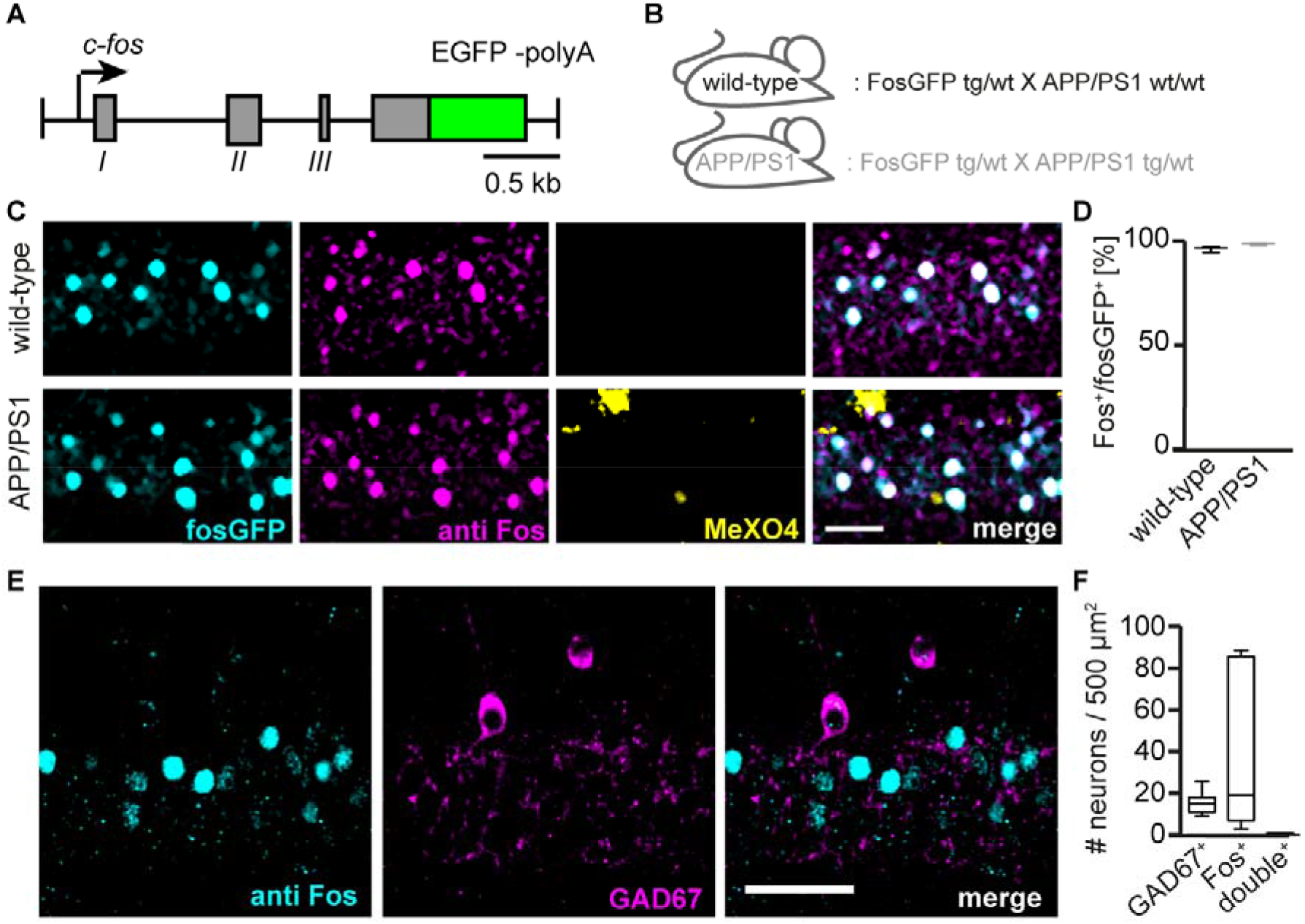
FosGFP reflects endogenous *c-fos* expression. **(A)** Schematic of the fosGFP transgene present in fosGFP mice. **(B)** FosGFP mice and APP/PS1 mice have been crossbred to analyze fosGFP expression in wild-type and APP/PS1 mice. **(C)** Exemplary confocal microscopy images of fosGFP-positive (fosGFP^+^) hippocampal CA1 neurons (cyan), double labeled with an anti Fos antibody (magenta) and the Aβ-plaque dye methoxy-X04 (MeX04, yellow). Wild-type and APP/PS1 mice are compared. **(D)** Percentage of fosGFP^+^ neurons double positive for endogenous Fos comparing wild-type and APP/PS1 mice. **(E)** Exemplary, fluorescence immunohistochemistry staining for Fos and GAD67 to label GABAergic neurons in hippocampal area CA1. **(F)** Box-plot diagram comparing the density of single and double positive GABAergic and Fos expressing neurons. (D) Data from n=3 wild-type, n=3 APP/PS1; p=0.052, unpaired t-test; (F) Data from n= 4 wild-type mice. Scale bar: (C) 40 µm, (E) 50 µm.

**Figure S2. Related to Figure 2.**
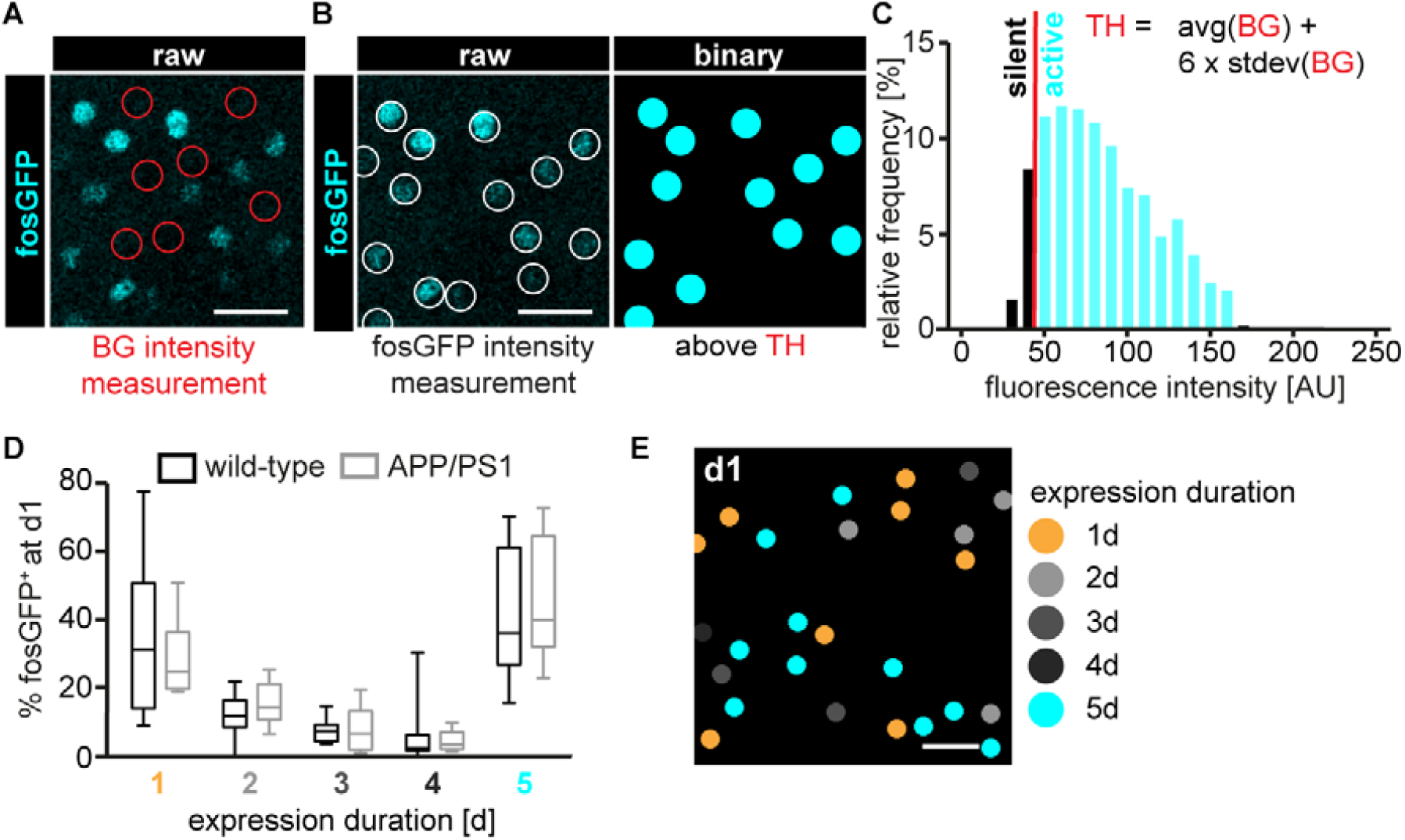
CA1 pyramidal neuron populations differ in their fosGFP expression dynamics. **(A-C)** Method to determine the threshold (TH) to generate binary images and categorize neurons as fosGFP^+^ (active) or fosGFP^-^ (silent). (A) Exemplary 8-bit fluorescence image and inserted circular masks (red) illustrating a measurement of background (BG) intensity. (B) Exemplary 8-bit fluorescence image before (raw) and after application of the threshold (binary). Inserted circular masks (white) illustrate a measurement of fosGFP intensity. (C) Exemplary fluorescence intensity histogram of fosGFP expressing cells with intensity below (silent, black) and above (active, cyan) threshold. The average (avg) BG intensity plus six fold its standard deviation (stdev) was used as threshold (TH). **(D)** Box-plot diagram comparing the fractions of neurons with different fosGFP expression durations between wild-type and APP/PS1 mice. **(E)** Exemplary binary image of fosGFP^+^ CA1 neurons color-coded according to their fosGFP expression duration. (D) Data from n=8 wild-type (4134 fosGFP^+^ neurons) and n=6 APP/PS1 mice (2993 fosGFP^+^ neurons) are compared; p=0.8966 (1d), p=0.9534 (2d), p=0.9630 (3d), p=0.9630 (4d), p=0.9534 (5d), two-way ANOVA with Holm-Sidak’s correction for multiple comparisons. Scale bars: 20 µm.

**Figure S3. Related to Figure 3.**
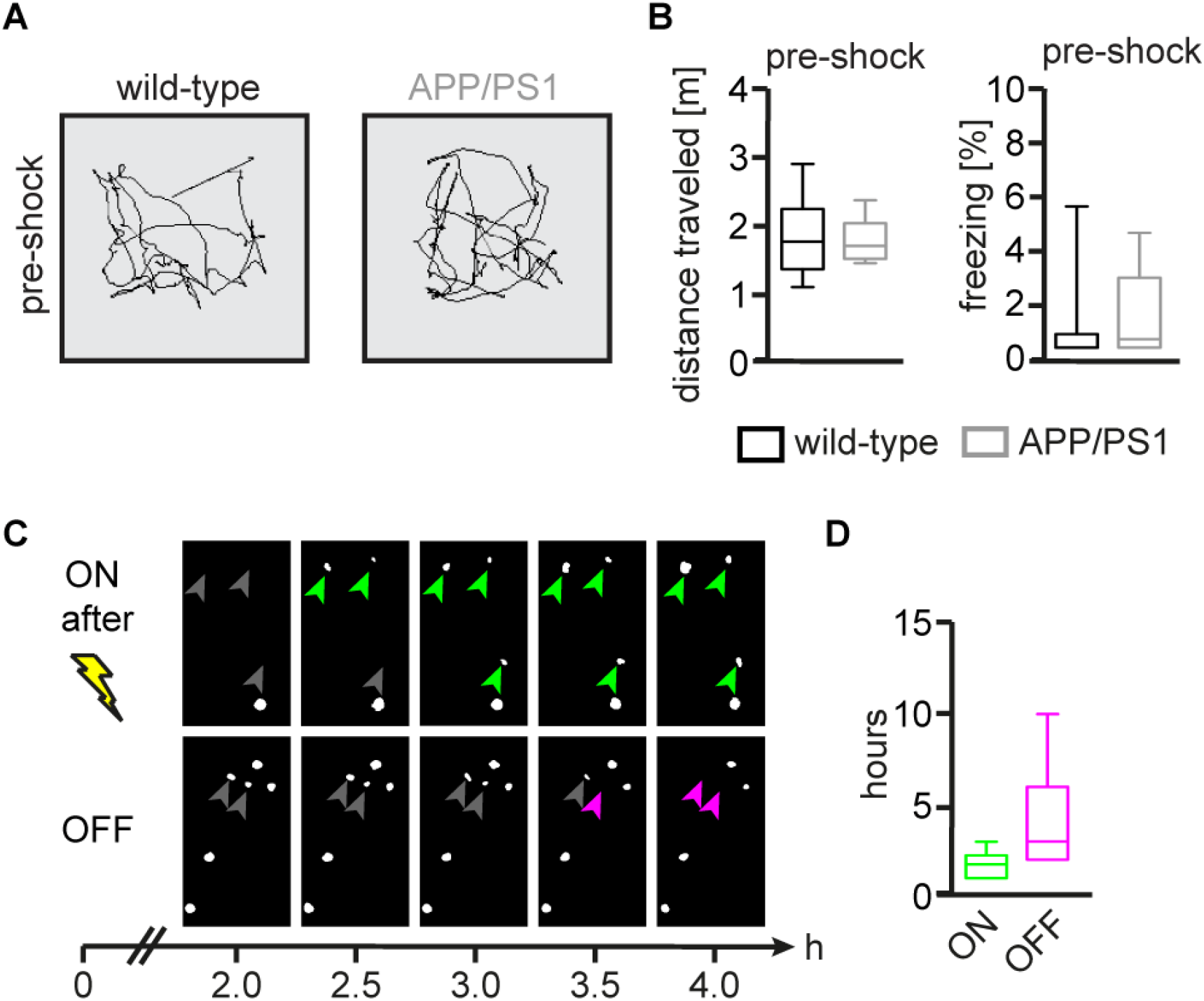
Additional behaviour parameters and fosGFP expression kinetics. **(A)** Exemplary traces of travelled distances during the one-minute interval before delivery of the first shock (pre-shock) comparing wild-type and APP/PS1 mice. **(B)** Average travelled distance (left) and freezing rate (right) of wild-type and APP/PS1 mice during the pre-shock interval. **(C)** Exemplary *in vivo* time-lapse images of fosGFP^+^ hippocampal CA1 neurons that turn “ON” (green arrows) or “OFF” (red arrows) fosGFP expression after contextual fear conditioning (yellow flash). **(D)** Average time to turn “ON” or “OFF” fosGFP expression. (B) Data from n=15 wild-type, n=11 APP/PS1; p=0.8514 (distance travelled, preFC), p=0.1213 (freezing, preFC), unpaired-t-test; (D) Data from n=52 fosGFP^+^ neurons.

**Figure S4. Related to Figure 3.**
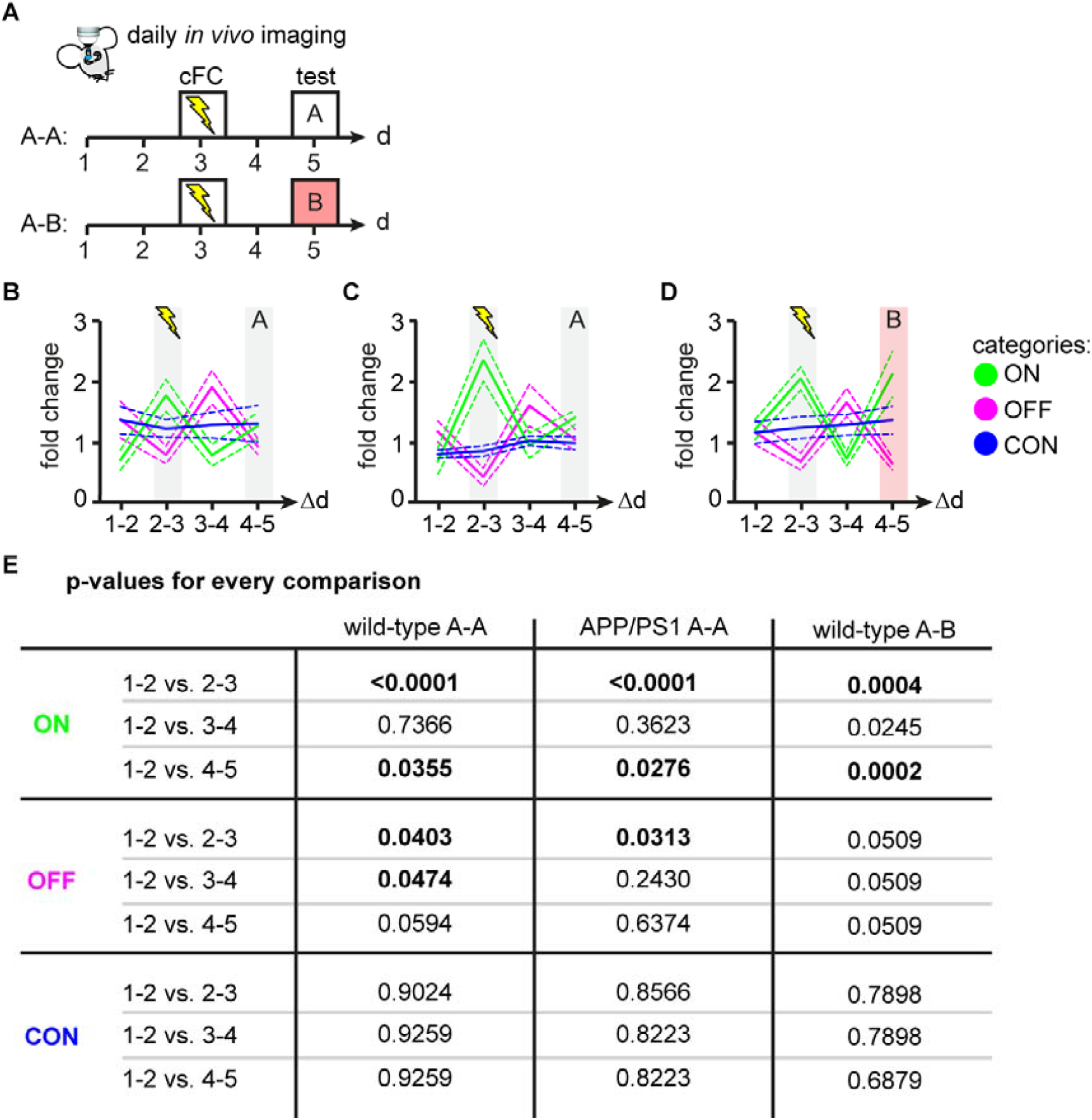
Intra-group comparisons of FosGFP expression dynamics during contextual fear conditioning and test. **(A)** Experimental timeline to investigate the dynamics of fosGFP expression after contextual fear conditioning (cFC) and memory test in the conditioned context A (A-A) or a novel context B (A-B). **(B-D)** Normalized fold changes of ON, OFF and CON neuronal categories in wild-type mice exposed to context A (B) or B (D), and (C) APP/PS1 mice exposed to the conditioned context A. **(E)** Individual p-values resulting from intra-group comparisons of ON, OFF and CON categories using a two-way ANOVA with Holm-Sidak’s correction for multiple comparisons. P-values <0.05 are displayed in bold. (B-E) Data from n=8 wild-type A-A (4775 neurons), n=6 APP/PS1 A-A mice (3776 neurons) and n=6 wild-type A-B mice (5099 neurons). (B-D) Data are presented as mean (continuous line) ± s.e.m. (dashed lines).

**Figure S5. Related to Figure 4.**
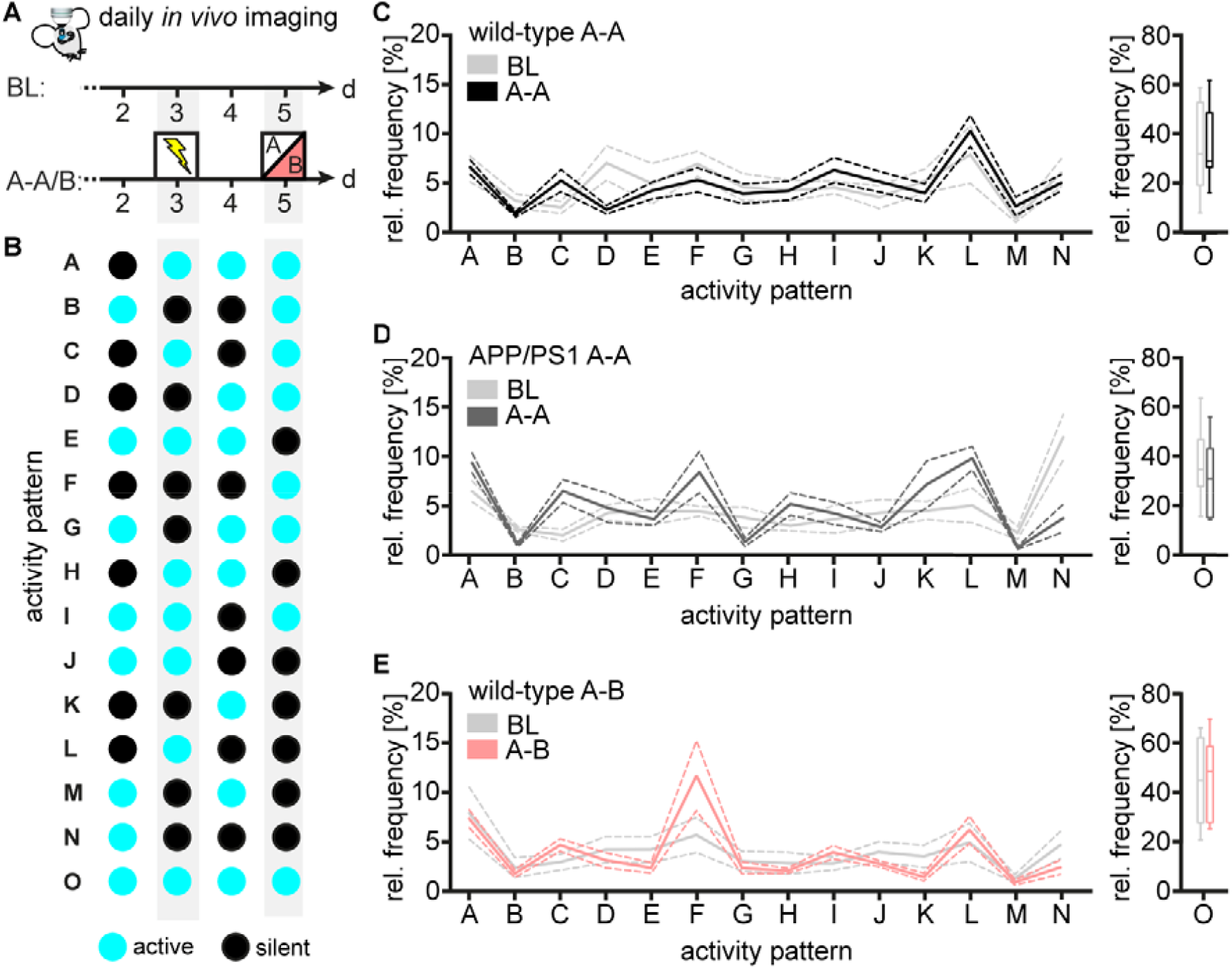
Relative activity pattern frequency of individual CA1 neurons. **(A)** Experimental time-line of *in vivo* fosGFP expression analysis to determine the activity state of CA1 neurons (active, silent) during baseline (BL) and contextual fear conditioning (A-A/B). Three experimental groups were compared: wild-type mice trained and tested in context A or B and APP/PS1 mice in context A. **(B)** Scheme illustrating every possible activity pattern (labeled from A to O) an individual neuron can adopt. Active and silent states are depicted in cyan and black, respectively. **(C-E)** Relative frequency of every possible activity pattern comparing BL and contextual fear conditioning (A-A/B) in wild-type A-A (C), APP/PS1 A-A (D) and wild-type A-B mice (E). Data from n=8 wild-type A-A mice (4775 fosGFP^+^ neurons), n=6 APP/PS1 A-A mice (3776 fosGFP^+^ neurons), n=6 wild-type AB mice (5099 fosGFP^+^ neurons). Data are presented in mean (continuous line) ± s.e.m. (dashed lines) (Pattern A-N) or in a boxplot diagram (Pattern O).

**Figure S6. Related to Figure 7.**
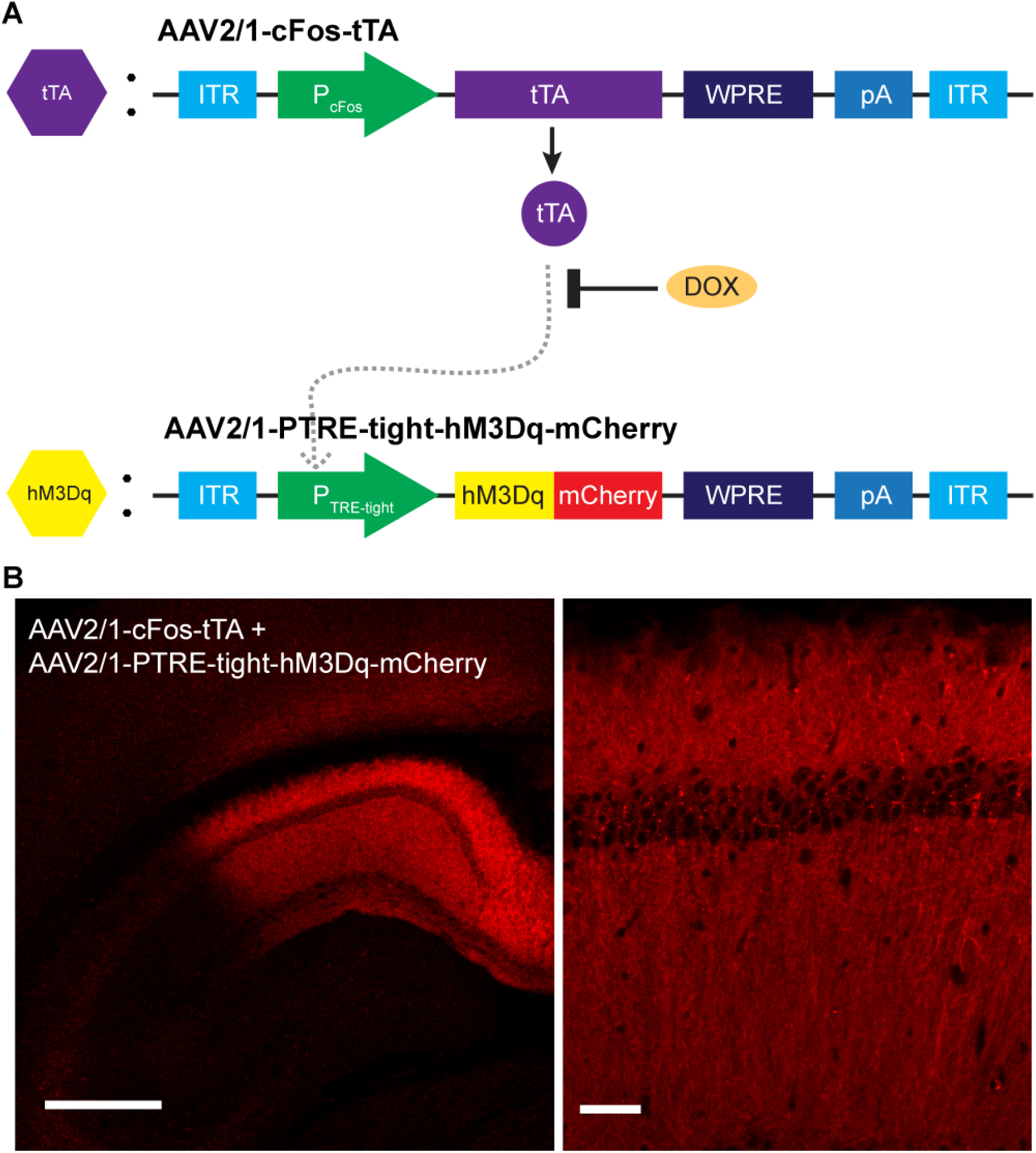
TetTag-system for activity-dependent labelling within a specified time window. **(A)** Scheme presenting the genomes of AAV2/1-sFos-tTA and AAV2/1-PTRE-tight-hM3Dq-mCherry. The transcriptional transactivator (tTA) is expressed in an activity-dependent manner utilizing the *c-fos* promoter (P_cFOS_). In the absence of Doxycycline (DOX) tTA binds to the promoter of hM3Dq (PTRE-tight) and starts its expression. Otherwise tTA is bound by DOX and hM2Dq expression is prevented. ITR, inverted double repeats; WPRE, woodchuck hepatitis virus posttranscriptional regulatory element; pA, polyadenylation site. **(B)** Overview (left) and enlarged (right) confocal images showing hM3Dq-mCherry expression in CA1. Scale bar: 500 µm (left), 50 µm (right).

